# Cell-autonomous restoration of splicing homeostasis and RP11 phenotype in patient-derived RPE and retinal organoids by *PRPF31*.AAV gene therapy

**DOI:** 10.64898/2026.04.14.718379

**Authors:** Maria Elia, Valda Pauzuolyte, Maria Georgiou, Mark Basche, Carina Hansohn, Elton J. R. Vasconcelos, Robert Atkinson, Sushma Grellscheid, Colin A Johnson, Gerrit Hilgen, Henning Urlaub, Alexander J. Smith, Sina Mozaffari-Jovin, Robin R. Ali, Majlinda Lako

**Affiliations:** Biosciences Institute, Faculty of Medical Sciences, Newcastle University, Newcastle upon Tyne, UK; Centre for Gene Therapy & Regenerative Medicine, Faculty of Life Sciences & Medicine, King’s College London, UK; Department of Clinical Chemistry, University Medical Center Göttingen, Göttingen, Germany; Max-Planck-Institute for Multidisciplinary Sciences, Göttingen, Germany; Leeds Omics, University of Leeds, UK; Department of Informatics, University of Bergen, Norway; Leeds Institute of Medical Research, Faculty of Medicine & Health, University of Leeds, UK; School of Geography and Natural Sciences, Northumbria University, United Kingdom, UK

**Keywords:** PRPF31, retinal organoids, gene therapy, retinal pigment epithelium cells, cilia, splicing, retinitis pigmentosa

## Abstract

Mutations in *PRPF31* gene cause retinitis pigmentosa type 11 (RP11) through haploinsufficiency, impairing spliceosome assembly and triggering progressive retinal degeneration. While gene augmentation holds therapeutic promise, key questions remain regarding the mechanistic basis of rescue and its therapeutic efficacy across all primarily affected human retinal cell types and disease stages. Here, we utilised patient-specific induced pluripotent stem cells (iPSCs)-derived retinal pigment epithelium (RPE) cells and three-dimensional retinal organoids (ROs) to determine the therapeutic mechanism of AAV-mediated *PRPF31* delivery. Using the ShH10(Y445F) serotype to ensure robust dual targeting of RPE and photoreceptors, we demonstrated that *PRPF31* transduction restores nuclear localisation, reorganises SC35^+^ nuclear speckles and enhances p-SF3B1⁺ active spliceosome foci. Transcriptomic and proteomic profiling revealed a global rescue of splicing activity and upregulation of phagocytosis, protein aggregate clearance pathways, and mitochondrial proteins. These molecular shifts facilitated the clearance of proteotoxic cytoplasmic aggregates and reversed key functional deficits; specifically, they reinforced RPE apical-basal polarity, restored phagocytic capacity and normal ciliary morphology and incidence, and boosted light-evoked activity in photoreceptors. Combining gene therapy with rapamycin-mediated-autophagy activation conferred no additive benefit, identifying the restoration of splicing homeostasis as the critical therapeutic driver. Notably, substantial phenotypic rescue is achievable in mature RPE, supporting a broad clinical window for intervention. Collectively, these data provide a systems-level validation of ShH10(Y445F)-PRPF31 gene therapy and establish a mechanistic framework for its clinical translation in RP11.

## Introduction

Retinitis pigmentosa (RP) is the most common form of inherited retinal blindness, with a prevalence of approximately 1 in 3,000–5,000 (1). It is a heterogeneous group of diseases characterised by the progressive degeneration of the mid-peripheral retina, leading to tunnel vision and blindness (2, 3). While Luxturna provides effective gene therapy for the specific *RPE65*-mutant subset (∼1–2% of RP cases), no approved treatments exist for the vast majority of RP genotypes. Mutations in over 80 genes cause non-syndromic RP, with pre-mRNA processing factor genes (*PRPF31*, *PRPF8*, *PRPF3*, *PRPF6*, *SNRNP200*) representing the second most common cause of the autosomal dominant form (4, 5). The PRPFs are components of the U4/U6.U5 tri-snRNP complex, a major subunit of the spliceosome that catalyses pre-mRNA splicing. Specifically, mutations in *PRPF31* are the most common, constituting 8.9% of autosomal dominant RP, leading to a type of RP known as RP11 (5, 6). Most *PRPF31* mutations generate null alleles that reduce total protein levels and impair spliceosome assembly; hence, RP11 development is widely attributed to haploinsufficiency (7–11). Although ubiquitously expressed, *PRPF* mutations predominantly cause retina-specific disease (12). Earlier studies suggested that retinal cells’ high alternative splicing rate renders them particularly sensitive to spliceosome disruption (13–15), providing a potential explanation for why mutations in ubiquitous splicing factors often lead to retina-specific phenotypes.

RP11 is characterised by primary rod photoreceptor degeneration followed by cone and RPE dysfunction. Mouse models show primary RPE changes and photoreceptor degeneration, but are limited by their short lifespan, failing to recapitulate the protracted, late-stage remodelling seen in human disease (5, 7, 16–18). Patient-specific modelling is now possible using induced pluripotent stem cell (iPSC)-derived RPE cells and retinal organoids (ROs) containing photoreceptors (19). Using this patient-specific model, we demonstrated that *PRPF31* mutations impair *in vivo* splicing specifically in patient-derived retinal cells, rather than in fibroblasts or iPSCs. Notably, this defect selectively disrupts splicing programmes that regulate RNA processing, creating a vicious cycle that exacerbates splicing deficiencies in retinal cells (20).

Our previous data showed that the mutant PRPF31 protein does not integrate into normal spliceosomal complexes but instead mislocalises to the cytoplasm, where it forms aggregates in photoreceptor and RPE cells (11). These cytoplasmic aggregates trap the wild-type PRPF31 protein, lowering its concentration in the nucleus. As a result, the formation of the tri-snRNP complex, an essential component of the spliceosome, is impaired, disrupting pre-mRNA splicing and contributing to retinal cell dysfunction. Both RP11-ROs and RPE cells exhibit abnormal nuclear speckle morphology, consistent with defective splicing factor reservoirs. At the cellular level, RP11-patient iPSC-derived RPE cells exhibit defective phagocytosis and abnormal ciliogenesis, which could be associated with the disruption of ciliary gene splicing downstream of the mutated *PRPF31* (8, 20). Similarly, RP11-ROs exhibit disrupted primary cilium morphology, photoreceptor degeneration, and impaired functional neural networks (20). Under normal conditions, the unfolded protein response (UPR) is triggered to prevent misfolded protein accumulation and aggregation. In RP11, however, both UPR and waste disposal mechanisms fail, leading to progressive buildup of insoluble aggregates containing key retinoid metabolism and visual cycle proteins, including RLBP1 (11).

Our previous studies established that the wild-type *PRPF31* expression is significantly reduced in RP11-RPE cells and ROs. Furthermore, the mutant protein fails to incorporate into spliceosome complexes, collectively resulting in a reduction of active spliceosomes (11, 20). These findings reinforce haploinsufficiency as the primary pathomechanism and identify *PRPF31* gene augmentation as a rational therapeutic strategy for RP11. Indeed, gene augmentation therapy using adeno-associated viral (AAV) vectors remains the leading approach for the treatment of inherited retinal diseases due to their ability to mediate long-term transgene expression in the retina, established safety profile and proven efficacy in several retinal therapies. Using this approach, an AAV2/Anc80 vector with a *CASI* promoter increased *PRPF31* expression and restored normal phagocytosis function and cilia length in RP11 iPSC-derived RPE cells (8). Similarly, an AAV2-PRPF31 with a modified capsid variant (AAV2-7m8) vector was used in RP11-ROs, where it prevented rod and cone photoreceptor degeneration (17). Despite the potential of AAV-mediated gene augmentation, a systematic investigation into its impact on the fundamental causative mechanisms of RP11 remains lacking. Specifically, the effects on PRPF31 localisation, nuclear speckle integrity, and the formation of cytoplasmic aggregates are yet to be explored. Notably, the field lacks evidence for the simultaneous therapeutic rescue of both RPE cells and ROs, a crucial step for the treatment of RP11.

In this study, we employed an AAV-mediated *PRPF31* gene augmentation strategy in patient iPSC-derived RP11-RPE cells and ROs to assess its therapeutic impact on RP11-associated phenotypes. We demonstrate that AAV-ShH10(Y445F)-mediated *PRPF31* delivery successfully restores both protein expression and physiological nuclear localisation in both RP-11 RPE cells and ROs. Furthermore, it leads to a significant increase in the abundance of active spliceosomes within RP11-photoreceptors and RPE cells, indicating a restoration of splicing machinery function, accompanied by improvement in transcriptomics and proteomics readouts. Functionally, we demonstrate that *PRPF31* gene augmentation improves the phagocytic function in RP11-RPE cells, increases cilia length and incidence, and inhibits cytoplasmic aggregate accumulation in both RPE cells and photoreceptors.

## Results

### Optimisation of AAV-Mediated Gene Delivery in iPSC-Derived RPE Cells and Retinal Organoids

Control and RP11-iPSCs were differentiated into RPE cells and ROs according to established protocols (11, 20). To determine the optimal dosage and transduction efficiency of *PRPF31*-AAVs in RPE cells, we performed optimisation studies using GFP-AAVs at single and double doses (administered one week apart) at MOIs of 0 to 1×10^6^ vg/cell across multiple control and RP11-RPE cells (**Figure S1A, B**). Statistical analysis revealed no significant difference in the percentage of transduced RPE cells between the single- and double-dose groups (**Figure S1C**). Furthermore, increasing the viral load beyond an MOI of 1×10^5^ vg/cell did not improve transduction rates in either treatment group (**Figure S1C**). Notably, cytotoxicity was significantly higher in groups receiving a double dose **(Figure S1D)**. Based on these findings, a single dose at an MOI of 1×10^5^ vg/cell was selected as the optimal condition, achieving high transduction efficiency (75.5% GFP+) while limiting cellular toxicity (**Figure S1E, F**).

Dose-response studies conducted across multiple control and RP11-iPSC-derived ROs, indicated a single dose at an MOI of 3 x 10^6^ vg/cell as the optimal condition. This condition resulted in a balanced transduction efficiency in CD73+ photoreceptors (17.5%; **Figure S2A–D, F, G**), while preserving cell viability and minimising toxic effects (**Figure S2E**).

### AAV-mediated PRPF31 Delivery Increases Nuclear PRPF31 Levels and Restores Nuclear Speckle Size in Retinal Cells

To evaluate the impact of *PRPF31* gene augmentation, we transduced RP11-RPE cells and ROs with either PRPF31.AAVs or a control vector (GFP.AAVs) using the optimal MOIs determined above. We then employed a combination of quantitative RT-PCR, western blotting, and immunofluorescence to evaluate the impact of *PRPF31* gene augmentation on the molecular and functional recovery of RP11-RPE cells and photoreceptors.

RP11-RPE cells were transduced three days post-plating with either PRPF31.AAVs or GFP.AAVs control at an MOI of 1 x 10^5^ MOI vg/cell. Two weeks post-transduction, qRT-PCR targeting the construct-specific 3’ UTR confirmed comparable AAV transgene expression levels in both groups (**Figure 1A**). While endogenous *PRPF31* expression levels remained unchanged across all groups, *PRPF31*-transduced cells exhibited robust transgene expression and a significant increase in total PRPF31 at both the mRNA and protein levels (**Figure 1A, B**).

**Figure 1:**
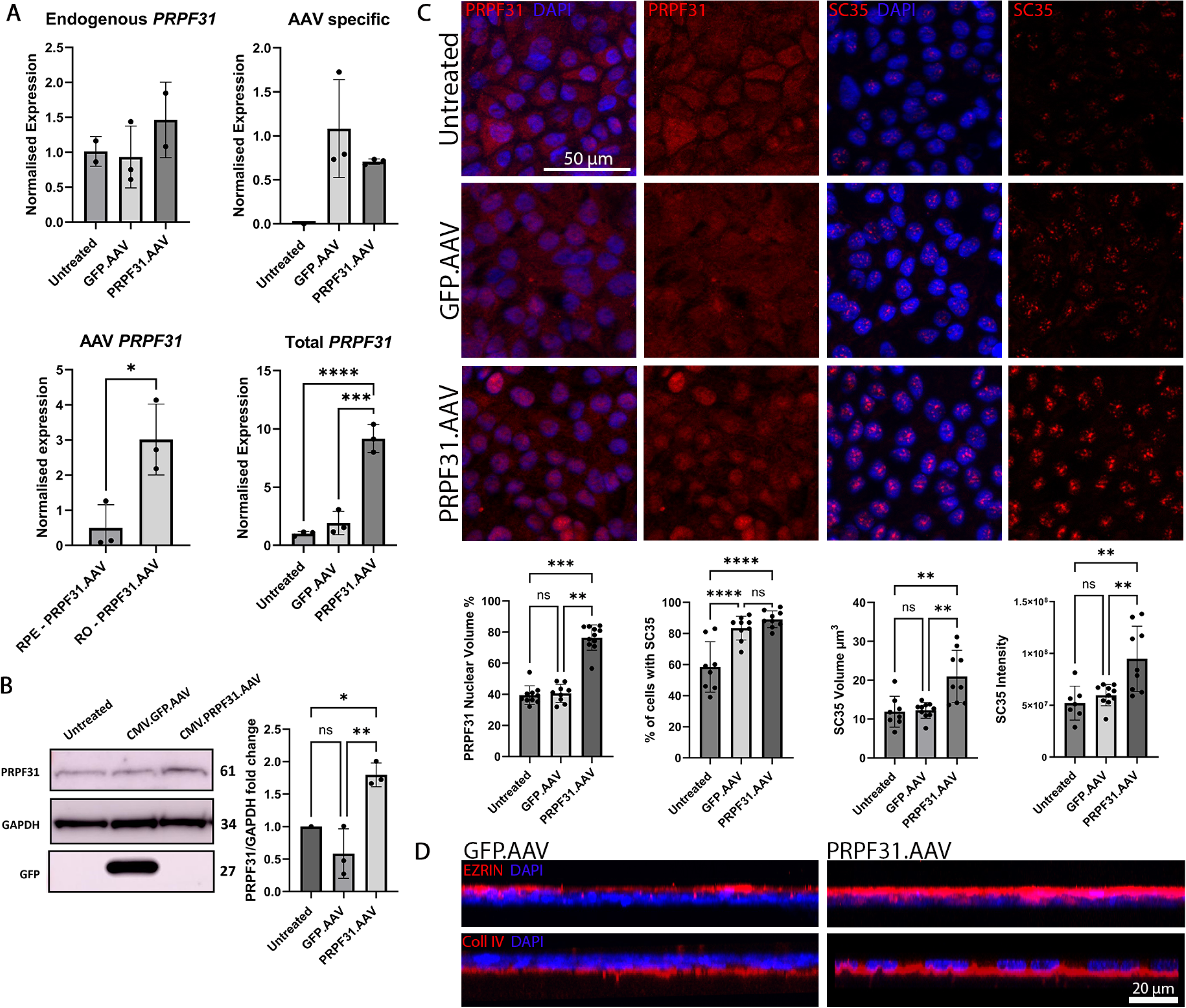
PRPF31.AAVs restore PRPF31 nuclear localisation, nuclear speckle organisation, and RPE cell polarity. **(A) Transcriptional upregulation of *PRPF31*.** Quantitative RT-PCR demonstrates significantly increased *PRPF31* mRNA levels in RP11-RPE cells following transduction with 1×10^5^ vg/cell *PRPF31*.AAV. Expression was normalised to *GAPDH*. **(B) PRPF31 protein recovery.** Representative Western blot and densitometric quantification confirm elevated PRPF31 protein expression in *PRPF31*. AAV-treated cells. Protein levels were normalised to GAPDH. **(C) Nuclear speckle reorganisation and PRPF31 localisation.** Immunostaining reveals the restoration of nuclear PRPF31 localisation (red) and reorganised SC35+ nuclear speckles (red) in treated RP11-RPE cells. Nuclei were counterstained with DAPI (blue). Quantitative analysis confirms significant increases in SC35+ speckle intensity, volume, and the total percentage of SC35+ cells. Scale bars: 50 µm. **(D) Reinforcement of apical-basal polarity.** Immunofluorescence imaging shows fully restored apical basal RPE cell polarity upon *PRPF31*.AAV treatment, evidenced by the continuous basal localisation of Collagen IV (Coll IV; red) and apical enrichment of EZRIN (red). Nuclei were counterstained with DAPI (blue). Scale bars: 20 µm. **Statistical Analysis:** Data are presented as mean ± SD. For (A) and (B), statistical significance was determined by one-way ANOVA with Tukey’s multiple comparisons test (n = 3 biological replicates). For (C), significance was assessed via Kruskal-Wallis’s test (PRPF31 and SC35 volume) or Tukey’s test (SC35 intensity and percentage); n = 9 representative images per marker per condition (*p < 0.05, **p < 0.01, ***p < 0.001, ****p < 0.0001).

We previously reported that in RP11-RPE cells, mutant PRPF31 is predominantly sequestered in the cytoplasm, forming insoluble aggregates that trap wild-type PRPF31, depleting it from the nucleus and impairing tri-snRNP assembly (11). Following PRPF31.AAV transduction, immunofluorescence microscopy revealed a marked increase in PRPF31 nuclear localisation (**Figure 1C**) with reduced cytoplasmic signal. This phenotype reflects two likely synergistic mechanisms: (i) AAV-driven expression of wild-type PRPF31 restores stoichiometric tri-snRNP assembly, promoting efficient nuclear import and retention; and (ii) re-establishment of normal spliceosomal function corrects the mis-splicing of autophagy and lysosome genes, activating autophagic clearance of pre-existing cytoplasmic aggregates. Together, these processes create a positive feedback loop that eliminates cytoplasmic PRPF31 sequestration and restores nuclear spliceosomal activity. The restored localisation pattern closely mirrored that observed in wild-type RPE cells (11); however, a residual cytoplasmic PRPF31 fraction persisted in AAV-transduced cells (23.5%) compared with a negligible cytoplasmic signal in wild-type controls (11). This indicates that while PRPF31 supplementation substantially rescues the pathological phenotype, it does not completely recapitulate the wild-type localisation profile.

Given that nuclear speckles have been identified as active hubs for co- and post-transcriptional pre-mRNA splicing, we next evaluated whether gene augmentation restored speckle morphology. Immunofluorescence analysis demonstrated a significant increase in the volume and fluorescence intensity of SC35^+^ nuclear speckles, alongside a higher percentage of SC35^+^ cells following treatment (**Figure 1C**). Beyond these molecular improvements, PRPF31.AAV administration reinforced apical–basal polarity, a critical physiological feature previously reported to be disrupted in RP11-derived RPE cells (20) (**Figure 1D**).

In RP11-ROs, *PRPF31* mutations also lead to the cytoplasmic mislocalisation of the PRPF31 protein and a concomitant reduction in nuclear speckle size within photoreceptors (11). To evaluate whether gene augmentation could reverse these defects, day 180 RP11-ROs were transduced with either PRPF31.AAVs or a GFP.AAVs. Two weeks post-transduction, qPCR and Western blot analyses confirmed a significant upregulation of both PRPF31 mRNA and protein levels in treated organoids (**Figure 2A, B**). Notably, the fold-increase in total PRPF31 protein expression was higher in ROs (3-fold) than in transduced RPE cells (1.8-fold).

**Figure 2:**
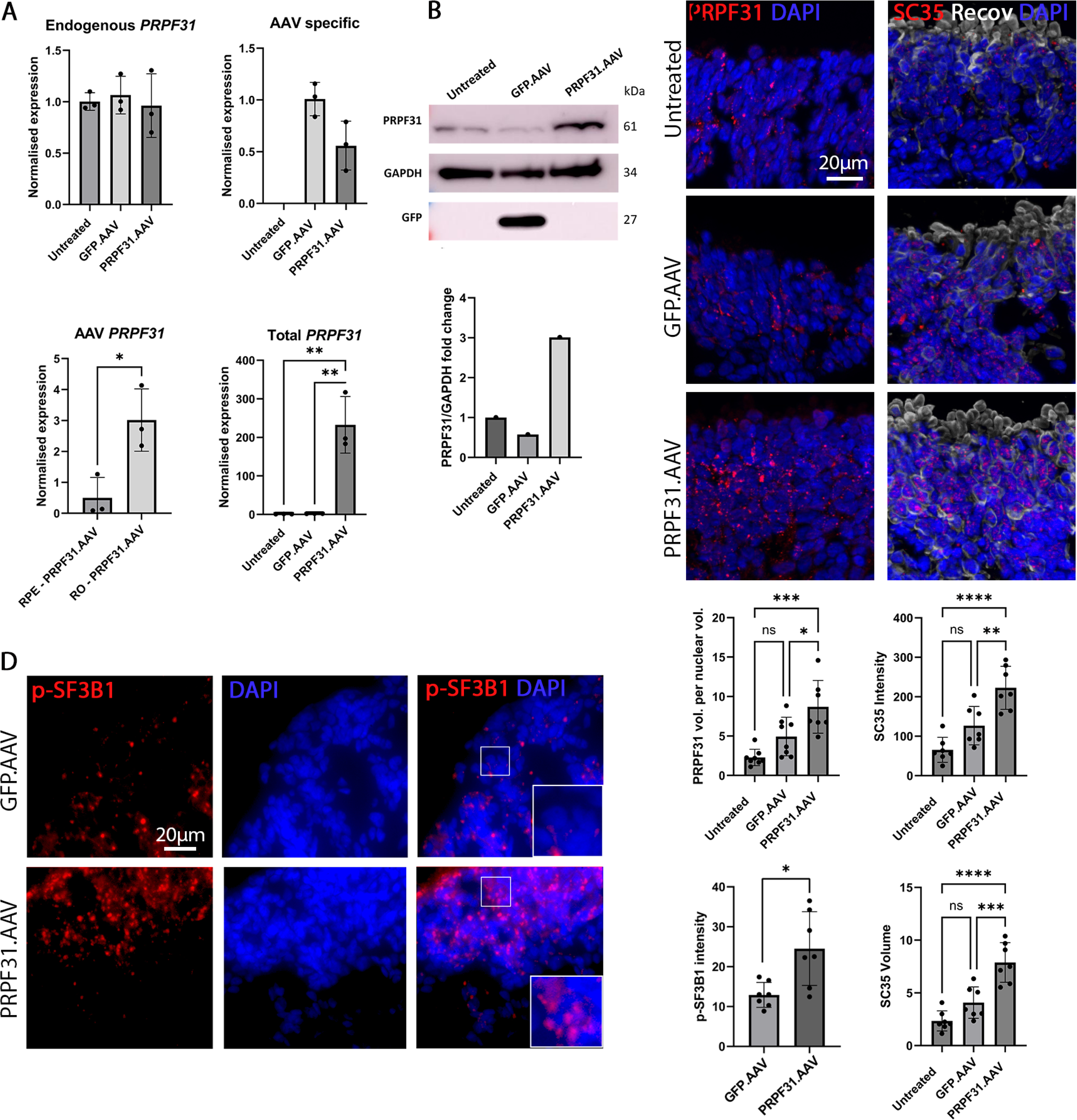
PRPF31.AAVs restore *PRPF31* expression, nuclear speckles, and the abundance of active spliceosomes in RP11-ROs. **(A) Quantitative RT-PCR analysis of *PRPF31* mRNA levels in RP11-ROs**. Transduction with 3×10^6^ vg/cell *PRPF31*.AAV significantly increases total *PRPF31* transcript expression compared to controls. Expression was normalised to *GAPDH*. **(B) Protein expression of PRPF31.** Representative Western blot (top) and densitometric quantification (bottom) demonstrate significantly elevated PRPF31 protein levels in *PRPF31*.AAV-treated RP11-ROs. Protein levels were normalised to GAPDH. **(C) Restoration of nuclear PRPF31 and SC35+ nuclear speckles architecture.** Immunostaining shows increased nuclear localisation of PRPF31 (red) and enhanced expression of the nuclear speckle marker SC35 (red) following treatment. Recoverin (white) labels photoreceptor outer segments; nuclei are counterstained with DAPI (blue). Quantitative analysis confirms significant increases in PRPF31 nuclear volume, as well as SC35 speckle intensity and volume, following PRPF31.AAVS treatment. Scale bar: 20 μm. **(D) Elevated abundance of active spliceosomes.** Immunostaining of RP11-ROs reveals increased staining of p-SF3B1 (red), a marker of active spliceosome (B^act^ and C) complexes, upon *PRPF31* augmentation. Nuclei are counterstained with DAPI (blue). Scale bar: 20 μm. **Statistical Analysis:** Data are presented as mean ± SD. Statistical significance was determined by one-way ANOVA followed by Tukey’s multiple comparisons test (*p < 0.05, **p < 0.01, ***p < 0.001, ****p < 0.0001). For (A) and (B), n = 3 biological replicates; for (C, D), n = 7 representative images per marker per condition.

This restoration of PRPF31 expression translated into structural improvements within the nucleus. We observed a significant increase in PRPF31 volume and intensity of SC35-stained nuclear speckles in PRPF31. AAV-treated ROs, suggesting a broad rescue of nuclear speckle integrity (**Figure 2C**). To determine if this structural rescue translated into functional recovery, we examined the formation of active spliceosomes via immunostaining for phosphorylated SF3B1 (p-SF3B1). As a component of the U2 snRNP, SF3B1 is specifically phosphorylated upon spliceosome activation (in B^act^ and C complexes), making it a robust indicator of splicing activity (21). RP11-ROs treated with PRPF31.AAV exhibited significantly more abundant and prominent p-SF3B1-enriched foci compared to controls (**Figure 2D**). These findings indicate that *PRPF31* augmentation successfully promotes spliceosomal assembly and enhances splicing activity within RP11 photoreceptors.

### PRPF31.AAV Treatment Prevents Cytotoxic Aggregate Accumulation and Rescues Ciliary Defects in PRPF31-Deficient RPE and Photoreceptor Cells

We previously demonstrated that mutant PRPF31 drives the progressive accumulation of cytotoxic cytoplasmic aggregates containing the visual cycle protein RLBP1 and the chaperone HSPB1 in retinal cells (11). To evaluate whether PRPF31 augmentation could mitigate this cellular pathology, we performed immunostaining for both markers two weeks post-transduction with PRPF31.AAV.

In RP11-RPE cells, treatment resulted in a marked reduction in the cytoplasmic abundance of RLBP1 and HSPB1 (**Figure 3A**), suggesting that PRPF31 expression promotes the clearance of these protein aggregates and/or prevents further proteotoxic aggregate accumulation. We observed a similar pattern of protein misfolding in ROs, where aggregate-like structures of HSPB1 and RLBP1 were detected between RP11-photoreceptors (**Figure 3B**). Consistent with our RPE findings, PRPF31 augmentation significantly decreased the prevalence of these inter-photoreceptor aggregates.

**Figure 3:**
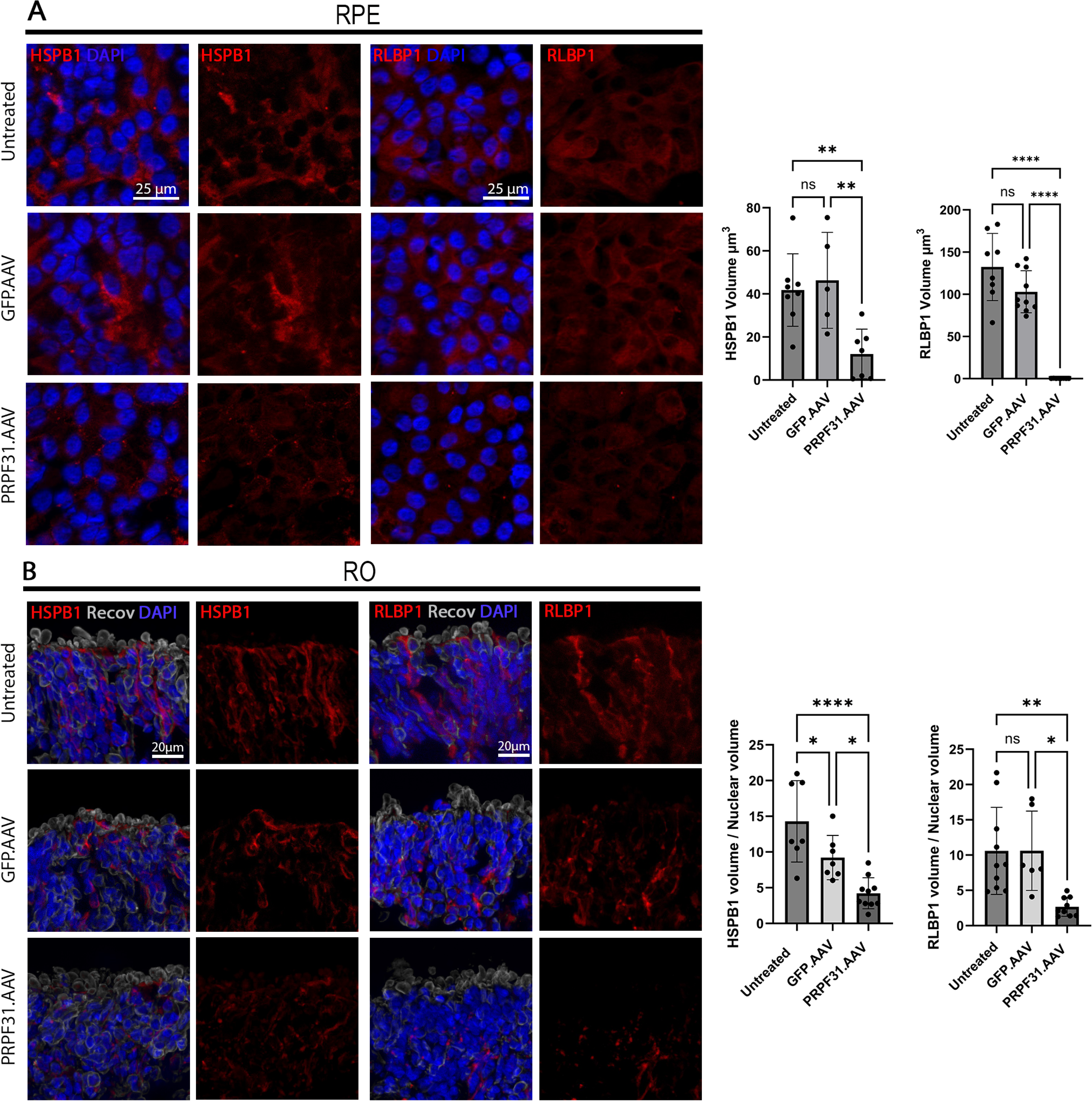
P*R*PF31 Augmentation Rescues Proteostasis by Clearing Cytoplasmic Aggregates in RPE and Retinal Organoids. **(A) Mitigation of protein aggregation in RP11-RPE cells.** Representative immunofluorescence images and volumetric quantification demonstrate a significant reduction in HSPB1 (red) and RLBP1 (red) cytoplasmic aggregates following PRPF31.AAV transduction. Nuclei were counterstained with DAPI (blue). Aggregate burden is expressed as total aggregate volume per cell. Scale bars: 25 µm. **(B) Resolution of aggregates in RP11-ROs.** Immunostaining of RO sections reveals that PRPF31.AAV treatment effectively clears HSPB1 (red) and RLBP1 (red) proteotoxic clusters within the photoreceptor layer, labelled with Recoverin (grey). Nuclei were counterstained with DAPI (blue). Quantitative analysis shows a significant decrease in aggregate volume normalised to total nuclear volume. Scale bars: 20 µm. **Statistical Analysis:** Data are presented as mean ± SD. Statistical significance was determined by one-way ANOVA followed by Tukey’s multiple comparisons test (*p < 0.05, **p < 0.01, ****p < 0.0001). For (A), n = 5 representative images per marker; for (B), n = 7 representative images per condition.

Beyond protein aggregation, we previously established that *PRPF31* mutations impair ciliogenesis, resulting in reduced cilia incidence and shortened cilia length in both RPE and photoreceptors (20). To assess the impact of PRPF31 gene therapy on cilia morphology, we immunostained PRPF31.AAV treated RP11-RPE cells and ROs for the ciliary marker ARL13B. Strikingly, AAV-mediated PRPF31 expression significantly increased both cilia length and incidence across both cell types (**Figure S3**). Thus, PRPF31.AAV gene therapy broadly reverses the cellular consequences of PRPF31 deficiency, addressing both protein aggregation and ciliary defects.

### PRPF31.AAV Delivery improves Phagocytic Activity in RPE cells and Enhances Light-evoked Responses in PRPF31-Deficient Photoreceptors

The phagocytosis of photoreceptor outer segments (POSs) is a hallmark of RPE health and a vital support function for retinal homeostasis. We previously reported that POS phagocytosis is severely impaired in RP11-RPE cells compared to wild-type controls (20). To determine whether *PRPF31* augmentation could restore this critical physiological process, we challenged RP11-RPE cells with pHrodo-labelled POSs and monitored internalisation over four hours via live-cell imaging. Our results demonstrate a robust functional improvement: PRPF31. AAV-treated RPE cells exhibited a nearly three-fold increase in POS internalisation (59.3% of total area) compared to GFP.AAV-treated controls (19.2%) (**Figure 4A**). Thus, gene augmentation not only corrects molecular deficiencies but also successfully reinstates the phagocytic capacity of RP11-RPE cells.

**Figure 4:**
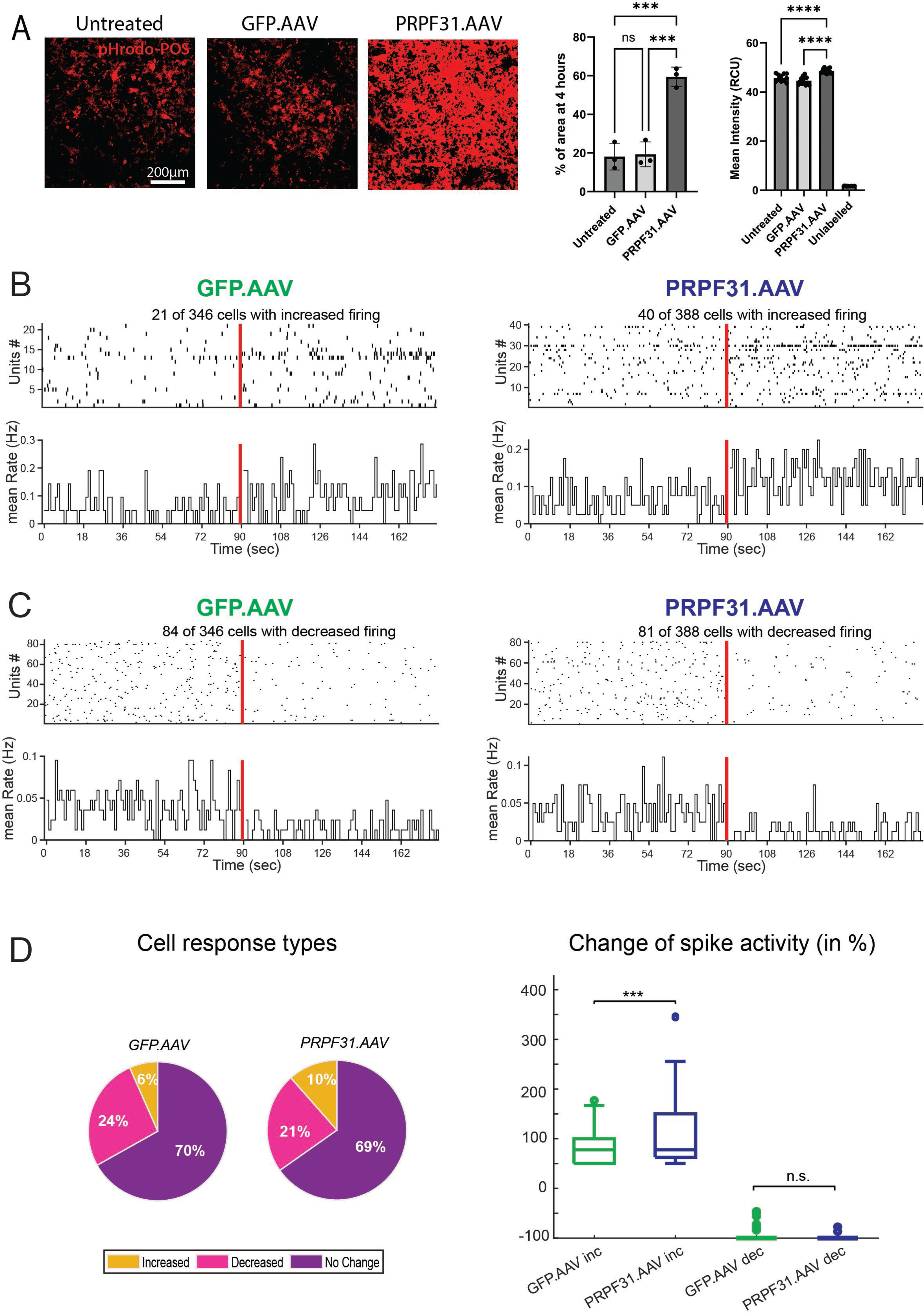
P*R*PF31 Augmentation Restores Phagocytic Capacity in RP11-RPE and Light-Evoked Circuit Activity in Retinal Organoids. **(A) Restoration of RPE phagocytosis.** RP11-RPE cells exhibit increased phagocytic capacity following PRPF31.AAV treatment. Internalised Photoreceptor Outer Segments (POSs) were labelled with pHrodo dye (red), which fluoresces upon acidification within the phagolysosome. Quantification displays the percentage of positive area and mean fluorescence intensity after 4 hours of incubation. Scale bars: 200 µm. **(B–C) Spatiotemporal RGC firing patterns.** Raster plots (top) and firing rate histograms (bottom; 5s bins) for 25 representative RGCs demonstrating either a 25% increase (B) or decrease (C) in spiking activity during pulsed light stimulation. In raster plots, vertical bars represent individual spikes; in histograms, the y-axis represents the mean spike rate across the population. The red line indicates the onset of pulsed light stimulation. **(D) Light-evoked RGC response distribution.** (Left) Heatmaps showing the population-wide distribution of RGC activity shifts in GFP.AAV (control) vs. PRPF31.AAV-treated ROs. Colour coding: yellow (no change), purple (= >50% increase), and pink (= > 50% decrease). (Right) Quantitative analysis of responsive RGC subpopulations. PRPF31.AAV treatment significantly increased the proportion of RGCs exhibiting “ON” (increased) responses compared to controls. No significant difference was observed in the “OFF” (decreased) subpopulation. **Statistical Analysis:** Data are presented as mean ± SD. For (A), significance was determined by one-way ANOVA with Tukey’s test (***p< 0.001, ****p < 0.0001), n=3-10 images per condition. For (D), an unpaired t-test was used to compare increased vs. decreased activity proportions (***p < 0.001). No significant difference was observed in decreased activity, n = GFP.AAV increased (21), PRPF31.AAV increased (40); GFP.AAV decreased (84), PRPF31.AAV decreased (81).

We previously demonstrated that RP11-derived ROs exhibit disrupted functional signalling, characterised by significantly diminished retinal ganglion cell (RGC) responses (20). In the current study, we assessed functional improvements by stimulating transduced ROs with different contrast light flashes. To quantify the spike activity patterns of RGCs transduced with GFP.AAVs and PRPF31.AAVs, we grouped RGCs based on whether they exhibited no change, an increase, or a decrease of 25% after light onset. **Figures 4B, C** show the spike activity (top plot, vertical lines) of individual RGCs (y-axis) over a selected amount of time (x-axis). These raster plots are accompanied by corresponding histograms that summarise activity, while the red line indicates the start of the light flashes. GFP. AAV-transduced RP11-ROs showed sparse or minimal spiking increase (**Figure 4B**: left panel) in response to light onset, while PRPF31.AAV-transduced ROs exhibited markedly enhanced light-evoked responses (**Figure 4B**: right panel). There were no differences between GFP.AAV- or PRPF31.AAV-transduced RO activity when assessing RGCs activity that was decreased (**Figure 4C**).

Next, we counted the number of RGCs in the three groups (**Figure 4D**: left panel) and calculated the median percentage change (**Figure 4D**: right panel) of RGCs with increased activity (GFP.AAV inc, PRPF31.AAV inc) and decreased activity (GFP.AAV dec, PRPF31.AAV dec). Although the proportion of light-responsive RGCs in the *PRPF31*.AAV group showed only modest shifts (a 4% increase in increased responses and a 3% decrease in decreased responses), these values remain within expected physiological variability (**Figure 4D**: left panel). Importantly, the magnitude of the increased response was significantly higher in *PRPF31*. AAV-transduced ROs (**Figure 4D**: right panel, PRPF31.AAV inc), suggesting reactivation of light-responsive photoreceptor networks.

The increased number of spikes and the stronger increased activity change imply that *PRPF31* augmentation improves synaptic transmission and downstream retinal response to light. These data suggest that *PRPF31* augmentation not only rescues individual cell function but also improves the overall functional integrity of the retinal circuitry.

### PRPF31 Augmentation Rescues Transcriptomic Defects in RP11 retinal Models

We previously reported that transcriptomic profiling of RP11-RPE cells and ROs revealed the most significant enrichment of Gene Ontology (GO) biological processes terms related to pre-mRNA and alternative splicing. These findings are consistent with the established role of *PRPF31* mutations in impairing spliceosomal assembly. Notably, the most differentially expressed genes (DEGs) in RP11-ROs were associated with the ciliary membrane and photoreceptor inner and outer segments, identifying cilia-related pathway defects as a primary driver of the retinal phenotype (20).

To characterise the molecular shifts induced by *PRPF31* augmentation, we performed bulk RNA-seq and proteomics on RP11-RPE cells and ROs treated with either PRPF31.AAVs or GFP.AAVs control. Differential expression analysis using DESeq2 identified fewer DEGs in RPE cells (n = 251) compared to the transcriptomic changes observed in ROs (n = 1128, **Table S1**). As expected, *PRPF31* was the most highly overexpressed gene in both treated models, showing log2 fold changes of 1.41 in RPE cells and 3.45 in ROs. To prevent this robust transgene expression from obscuring more subtle downstream transcriptomic shifts, *PRPF31* was excluded from subsequent pathway and enrichment analyses.

In RP11-RPE cells, RNA-seq analysis revealed that gene sets associated with amyloid-beta clearance pathways were significantly upregulated following treatment (**Figure 5A**). This transcriptomic shift aligns with the observed reduction in cytotoxic protein aggregates (**Figure 3A**). Specifically, the DEGs driving this enrichment included *HMGCR, LDLR, LRP2,* and *CYP51A1,* key players in cholesterol metabolism and lipoprotein receptor-mediated endocytosis. Conversely, genes involved in pro-inflammatory signalling pathways, particularly those mediated by TNF and IL-17, were enriched among the downregulated transcripts (**Figure 5A**). These results suggest that *PRPF31* augmentation may mitigate the chronic inflammation often associated with retinal degeneration in RP11.

**Figure 5:**
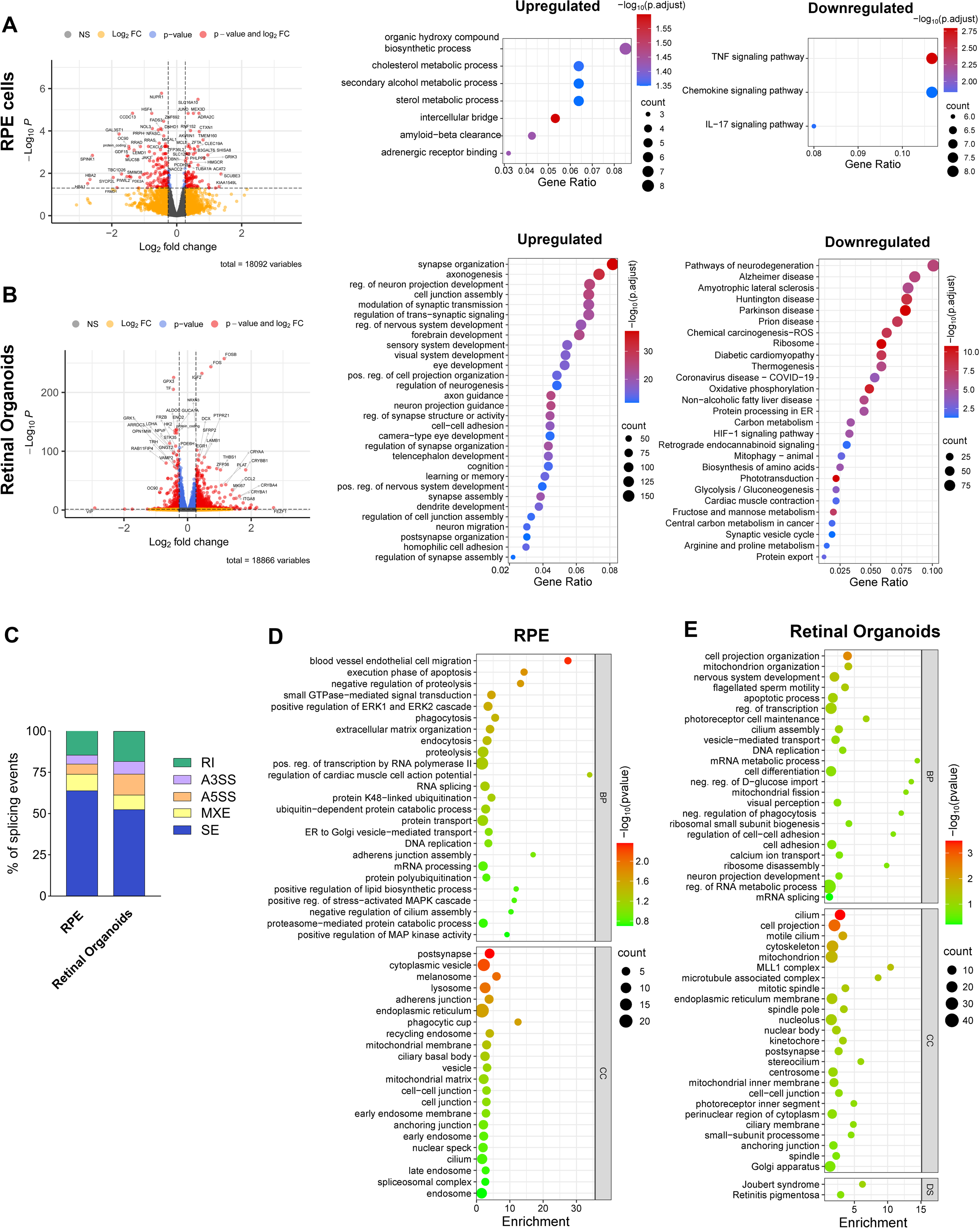
P*R*PF31 *Augmentation* Drives Global Transcriptomic Recovery and Alternative Splicing Modulation. **(A)** Differential gene expression in RP11-RPE cells. Volcano plot (left) identifies 251 differentially expressed genes following PRPF31.AAV treatment compared to GFP controls. Significance thresholds: padj-value < 0.05 and fold change FC ≥ 1.20 for upregulated genes or ≤ 0.83 for downregulated genes. Gene Ontology (GO) enrichment analysis (right) highlights restored biological pathways, including key metabolic and homeostatic signatures. **(B)** Transcriptomic shifts in RP11-ROs. Volcano plot (left) reveals 1128 DEGs modulated by PRPF31 overexpression. Significance thresholds: padj-value < 0.05 and fold change FC ≥ 1.20 for upregulated genes or ≤ 0.83 for downregulated genes. GO enrichment analysis (right) identifies significant modulation of pathways critical to neuroretinal function and survival. **(C)** Bar chart showing the percentage of differential alternative spliced events in PRPF31.AAV-treated ROs and RPE cells. rMATS analysis indicated that RP11-ROs have the highest proportion of transcripts containing alternative 3′ splice sites (A3SS). **(D)** Functional enrichment of differential alternative splicing in RPE. DAVID enrichment analysis of differentially spliced transcripts (identified via rMATS) in treated RP11-RPE cells. Significant enrichment is observed in biological processes related to mRNA splicing, spliceosomal assembly, cell adhesion, protein ubiquitination, and intracellular signalling cascades. **(E)** Gene ontology enrichment of differential splicing in ROs. DAVID analysis of differentially spliced transcripts in *PRPF31*.AAV-treated ROs. Cellular Component (CC) terms highlight significant association with the cilium, motile cilium, and photoreceptor inner segment, indicating structural restoration at the mRNA isoform level.

RNA-seq analysis of PRPF31.AAV-treated RP11-ROs revealed a significant upregulation of gene sets involved in synaptic function and visual system development, aligning with the improved retinal circuitry integrity shown above. In contrast, there was a marked downregulation of transcripts associated with neurodegenerative pathologies characterised by protein aggregation and RNA metabolism dysfunction, such as Alzheimer’s disease and Amyotrophic Lateral Sclerosis (**Figure 5B**).

We next investigated the impact of *PRPF31* augmentation on RNA splicing by examining differential exon usage across RPE cells and ROs. Utilising rMATS software (threshold values: p<0.05 and inclusion difference >0.05), we categorised alternative splicing events into five distinct types: skipped exons, retained introns, alternative 5’ and 3’ splice sites, and mutually exclusive exons. Our analysis revealed a comparable burden of differential splicing events in both tissues following *PRPF31*.AAV treatment, with 241 significant events identified in RPE cells and 261 in ROs (**Table S2**). ROs were characterised by a higher level of transcripts with alternative 3’ splice sites, while skipped exons were the most abundant splicing event in both ROs and RPE cells (**Figure 5C**).

To determine the functional consequences of these transcriptomic shifts, we performed gene enrichment analysis on the affected genes. In *PRPF31*.AAV-treated RPE cells, differentially spliced transcripts were significantly enriched for biological processes central to RNA metabolism, including mRNA splicing via the spliceosome and mRNA processing. Enrichment was also observed for phagocytosis, proteolysis, cell adhesion, and regulation of cilium assembly (**Figure 5D; Table S2**). These findings suggest that *PRPF31* augmentation directly modulates the splicing machinery itself, potentially initiating a compensatory feedback loop that restores homeostatic splicing patterns within the retinal environment. This molecular rescue likely underpins the reversal of structural and functional deficits observed in RP11-RPE cells, including improved phagocytosis, reduced aggregate accumulation, and ciliogenesis.

In parallel with our RPE findings, Gene Ontology (GO) enrichment analysis of differentially spliced transcripts in *PRPF31*.AAV-treated ROs revealed significant modulation of key regulatory networks. Specifically, these transcripts were enriched for biological processes critical to retinal homeostasis, including cilium assembly, transcriptional regulation, and regulation of RNA metabolic process, including mRNA splicing (**Figure 5E; Table S2**). Moreover, key pathways related to apoptosis, cell adhesion and mitochondria are enriched. To pinpoint where these transcriptomic changes manifest structurally, we performed Cellular Component (CC) GO analysis. This revealed that the alternatively spliced transcripts in *PRPF31*.AAV-treated RP11-ROs were predominantly associated with the cilium, synapses, mitochondria, and notably, the photoreceptor inner segment (**Figure 5E; Table S2**). This localised enrichment pattern is particularly significant, as the connecting cilium and inner segment are the primary sites of protein transport, metabolic activity, and energy production in photoreceptors. The restoration of splicing patterns in transcripts associated with these specific compartments suggests that *PRPF31* augmentation may directly restore the structural integrity, metabolic function, and trafficking capabilities of the RP11-affected retina.

### AAV-Mediated PRPF31 Augmentation Reshapes the Proteomic Landscape of RP11-RPE Cells and Photoreceptors Restoring Metabolic and Visual Cycle Pathways

To further evaluate the impact of *PRPF31* overexpression at the protein level, we performed quantitative proteomic analysis on RP11-RPE cells and ROs using Tandem Mass Tag (TMT) labelling followed by mass spectrometry (**Tables S3, S4**). Consistent with our transcriptomic findings, proteomics confirmed a significant upregulation of the PRPF31 protein in both models, following gene augmentation, validating the efficiency of the transgene expression (**Figure 6A, B**).

**Figure 6:**
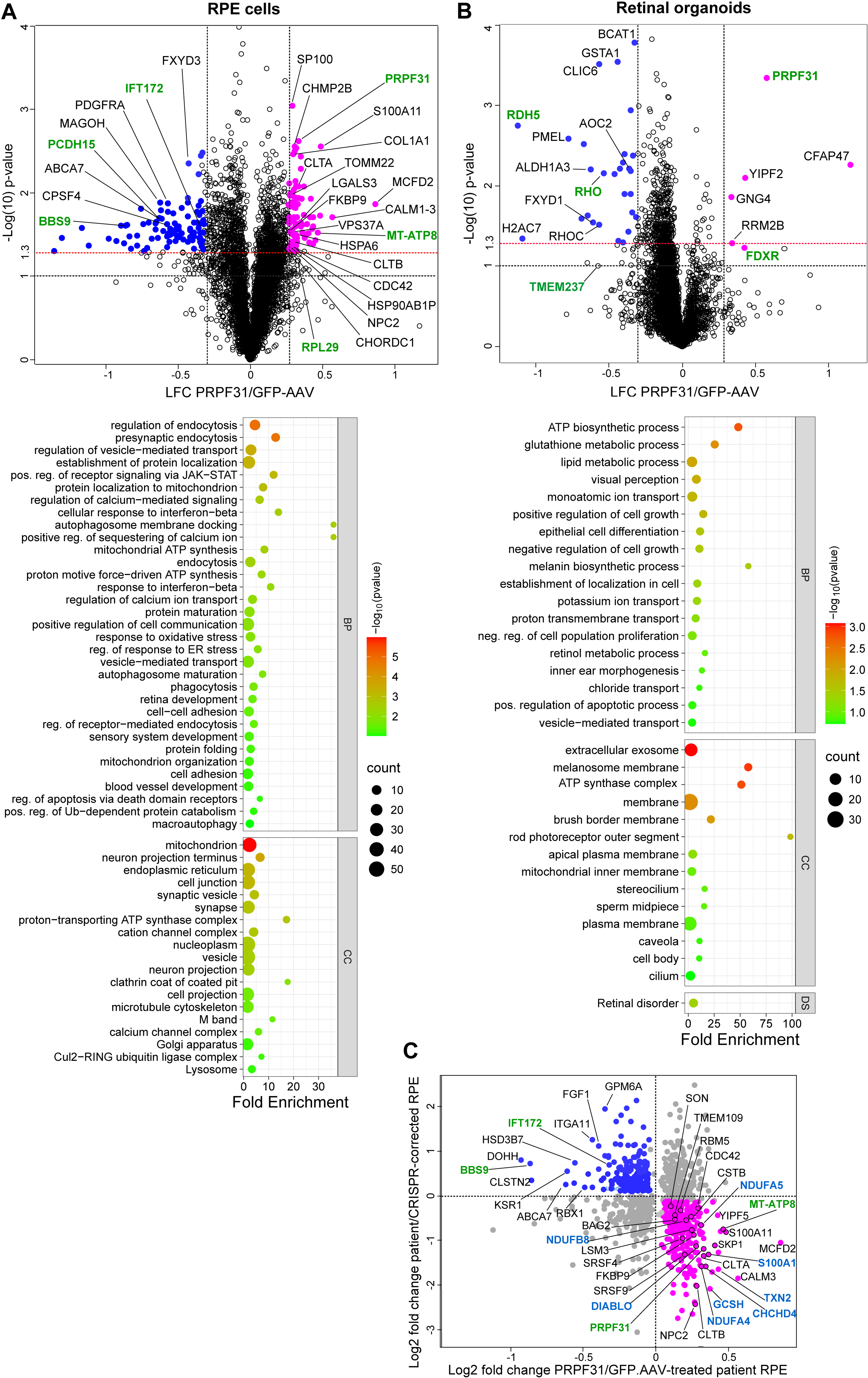
P*R*PF31 Augmentation Rescues Proteomic Signatures Linked to the Visual Cycle and Metabolic Homeostasis. **(A)** Proteomic landscape of RP11-RPE cells. Volcano plot (left) identifies 230 differentially expressed proteins following PRPF31.AAV treatment. Significance thresholds: padj-value < 0.05; log2 fold change (LFC) ≤ -0.26 for downregulated proteins; upregulated proteins LFC ≥ 0.26. Proteins previously implicated in the pathogenesis of Retinitis Pigmentosa (RP) are highlighted in green. Functional enrichment analysis (right) reveals that PRPF31 overexpression significantly modulates biological and cellular processes associated with RPE metabolic health and protein trafficking. **(B)** Proteomic shifts in RP11-ROs. Volcano plot (left) illustrates 46 differentially expressed proteins in PRPF31.AAV-transduced ROs. Significance thresholds: padj-value < 0.05; LFC ≤ -0.26 for downregulated proteins; upregulated proteins LFC ≥ 0.26. RP-associated proteins are indicated in green. Gene ontology enrichment analysis (right) identifies significant shifts in biological pathways critical for photoreceptor maintenance, including the visual cycle and ciliary transport. **(C)** Quadrant scatter plot correlating log2 fold changes of proteins in PRPF31. AAV-treated versus GFP. AAV-treated patient RPE with log2 fold changes in patient RPE versus its isogenic CRISPR-corrected cell line. Proteins downregulated in patient RPE and upregulated following PRPF31.AAV transduction are shown in pink, and those upregulated in patient RPE and downregulated in PRPF31.AAV transduced RPE are shown in blue. Key mitochondrial proteins and RP-associated proteins significantly regulated are highlighted in light blue and green, respectively.

Our analysis demonstrated that the proteomic landscape of the RPE cells was more extensively altered than that of the ROs. We identified significantly fewer differentially expressed proteins (DEPs) in treated ROs (n = 46) compared to the more profound proteomic shift observed in the RPE cells (n = 230). These findings indicate that although *PRPF31* therapy targets common pathways in both tissues, the RPE exhibits a more comprehensive proteomic shift. This suggests that the RPE may experience greater functional impairment in RP11, necessitating a more extensive cellular response to gene augmentation.

GO enrichment analysis of biological processes revealed that mitochondrial ATP synthesis, endocytosis, cell adhesion, protein folding, phagocytosis and macroautophagy were among the most significantly altered pathways in PRPF31.AAV-transduced RPE cells. Specifically, we observed increased expression of key proteins essential for endocytosis and phagocytosis, including CLTA, CLTB, G2MA, NPC2, LGALS3, and CDC42—the latter an established regulator of POS phagocytosis by RPE cells (22) (**Table S3**). These pathways are critical for RPE function, as they support the high metabolic demands of the retina and facilitate the phagocytosis of POS, a physiological process we demonstrated to be restored following *PRPF31* augmentation. Additionally, several proteins involved in autophagy—a pathway critical for clearing protein aggregates and damaged organelles (e.g. mitophagy)—were upregulated, including CALM3, CHMP2B, VPS37A, and NPC2.

Consistent with these findings, cellular component analysis identified the mitochondria, endoplasmic reticulum, and nucleoplasm as the most affected compartments (**Figure 6A**). The enrichment of nucleoplasmic proteins, which facilitate the diffusion of molecules required for DNA replication, transcription, and RNA processing, underscores a broader recovery of RPE homeostasis.

Given that our previous proteomic study and an RP11 mouse model identified RPE as the most affected cell type in RP11(20, 23), we next compared our PRPF31.AAV proteomic data with those from CRISPR-corrected patient iPSC-derived RPE (**Figure 6C**). Notably, 118 proteins from our list of DEPs in the PRPF31.AAV-treated RPE were also differentially expressed following CRISPR genome editing, with 48 proteins upregulated and 28 proteins downregulated in both conditions. Among these, several key mitochondrial proteins were substantially upregulated (coloured blue in **Figure 6C**). For example, MT-ATP8, a subunit of mitochondrial ATPase whose variants are a primary cause of NARP syndrome, where RP is a clinical feature (24), was significantly increased. Similarly, TXN2, a mitochondrial protein component of the thioredoxin system that regulates redox balance, was upregulated (25); its overexpression protects RPE against oxidative stress. Notably, key regulators of alternative mRNA splicing, including SRSF4, SRSF9, and RBM5 were also among the proteins upregulated after CRISPR gene editing or PRPF31 augmentation. RBM5 acts as a spliceosome gatekeeper at early stages of assembly and controls apoptosis (26). Altogether, this proteomic profile suggests that *PRPF31* augmentation rewires the RP11 cell proteome toward a healthy state, thereby restoring the cell’s capacity for POS phagocytosis and autophagy, and potentially contributing to the clearance of cytotoxic aggregates and damaged mitochondria observed in this study.

Proteomic analysis of PRPF31.AAV-treated RP11-ROs revealed significant alterations in biological processes related to ATP biosynthesis, lipid and retinol metabolism, the visual cycle, and apoptotic pathways. Cellular component enrichment identified proteins associated with rod outer segments (e.g., RHO, TMEM237), mitochondria and cilia. Within the visual cycle and retinol metabolism pathways, several proteins typically dysregulated in RP, including RHO, RDH5, were modulated following *PRPF31* augmentation (**Figure 6B, Table S4**). In addition, key proteins critical for mitochondrial function were substantially upregulated. These included RRM2B, whose variants are associated with mtDNA depletion and subsequent photoreceptor degeneration (27), and FDXR, a mitochondrial membrane protein essential for electron transport; mutations in *FDXR* lead to photoreceptor loss and a form of RP (28). Taken together, our transcriptomic and proteomic analyses demonstrate that *PRPF31* gene augmentation not only restores PRPF31 protein levels but also fundamentally reshapes the cellular and metabolic landscape of both RP11-RPE cells and ROs, driving them toward a healthy, functional state.

### *PRPF31* Augmentation Rescues Late-Stage RP11-RPE Pathology

Prolonged culture of iPSC-derived RPE cells promotes cellular maturation and induces ageing-associated features, an approach valuable for modelling late-stage retinal degeneration diseases (29). Mature RPE cell models often exhibit hallmarks of age-related pathophysiology, including accumulation of intracellular debris and formation of drusen-like deposits, as reported in long-term RPE culture models of age-related macular degeneration (AMD) (30, 31). Our previous work established that RP11-RPE cells progressively accumulate cytoplasmic aggregates, a process driven by a compromised autophagic degradation pathway. Notably, AAV-mediated gene augmentation has proven effective in rescuing key functional deficits in mature RP11-iPSC-RPE cells, specifically restoring ciliogenesis and partially recovering epithelial barrier function (8).

To determine whether *PRPF31* gene augmentation remains effective in late-stage disease, we transduced mature RP11-RPE cells (cultured on transwells for >12 weeks) with either GFP.AAVs or PRPF31.AAVs for two weeks. We then performed immunofluorescence analysis targeting PRPF31, nuclear speckles, cytoplasmic aggregates, and cilia, markers previously shown to be rescued by *PRPF31* augmentation in earlier disease stages. PRPF31.AAV transduction induced a significant reduction in PRPF31 cytoplasmic localisation compared to GFP.AAV transduced and untreated RPE cells, in accordance with the activation of macroautophagy observed in our proteomic analysis and restoration of nuclear speckles. Furthermore, *PRPF31* augmentation in these mature cells led to a measurable increase in both the volume and fluorescence intensity of SC35+ nuclear speckles (**Figure 7A**). These results align with our observations in early-stage RP11-RPE cells, confirming that *PRPF31* gene augmentation remains effective even in mature, late-stage models.

**Figure 7:**
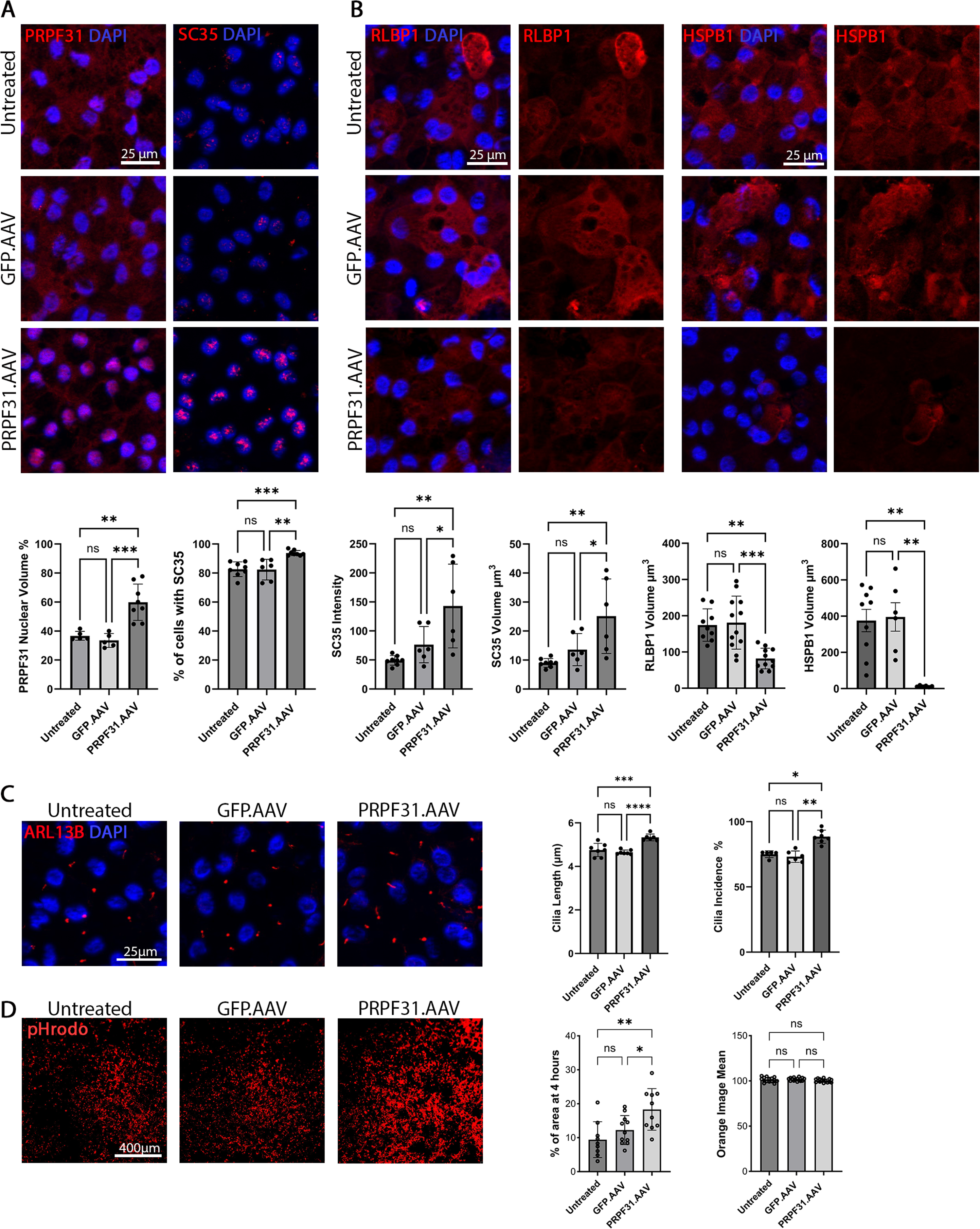
Late-Stage PRPF31 Augmentation Reverses Established Cellular Phenotypes in Mature RP11-RPE Cells. **(A)** Restoration of nuclear homeostasis. Mature RP11-RPE cells (treated post-maturation) show restored nuclear localisation of PRPF31 (red) and reorganised SC35+ nuclear speckles (red). Quantitative analysis confirms significant increases in SC35+ area, fluorescence intensity, and the percentage of SC35+ cells following *PRPF31*.AAV treatment. Nuclei are counterstained with DAPI (blue). Scale bars: 25 µm. **(B)** Clearance of established proteotoxic aggregates. Immunostaining reveals a significant reduction in the volume of HSPB1 (red) and RLBP1 (red) cytoplasmic aggregates in mature RPE cells upon treatment. Aggregate burden is expressed as the total aggregate volume per cell. Nuclei are counterstained with DAPI (blue). Scale bars: 25 µm. **(C)** Rescue of ciliary architecture. Immunofluorescence analysis using the ciliary marker ARL13B (red) demonstrates that *PRPF31* augmentation restores both cilia length and cilia incidence (percentage of ciliated cells) in mature RP11-RPE, reversing the stunted ciliary phenotype observed in controls. Nuclei are counterstained with DAPI (blue). Scale bars: 25 µm. **(D)** Recovery of functional phagocytosis. Phagocytic capacity in mature RPE is significantly improved following treatment, as evidenced by the internalisation of pHRodo-labeled Photoreceptor Outer Segments (POS; red). Data are quantified by positive area percentage and mean fluorescence intensity after 4 hours of incubation. Statistical Analysis: Data are presented as mean ± SD. Statistical significance was determined by one-way ANOVA with Tukey’s test, except for cilia length (C), which was assessed via Kruskal-Wallis with Dunn’s multiple comparisons test. Sample sizes: (A) n =5, (B) n=6, (C) n=7, (D) n=3-10, representative images per marker/condition. (*p < 0.05, **p < 0.01, ***p < 0.001, ****p < 0.0001).

Next, we investigated whether PRPF31 gene augmentation could eliminate the accumulation of cytoplasmic aggregates in mature RP11-RPE cells. Following transduction with PRPF31.AAV, immunostaining for RLBP1 and HSPB1 revealed a significant reduction in aggregate formation, mirroring the rescue observed in our earlier analyses (**Figure 7B**). Furthermore, PRPF31 augmentation significantly increased both cilia length and incidence compared to GFP.AAV controls (**Figure 7C**). To evaluate the functional impact of this treatment, we assessed phagocytic activity two weeks post-transduction using pHRodo-labeled POSs. PRPF31.AAV treatment resulted in a nearly two-fold increase in POS internalisation (18.33%) compared to untreated and GFP.AAV-treated controls (9.42% and 12.25%, respectively) (**Figure 7D**).

Collectively, these findings demonstrate that PRPF31.AAV effectively restores nuclear localisation and speckle formation, reduces proteostatic stress, and enhances phagocytic function in mature cells. Notably, however, the therapeutic effect was less pronounced than in early-stage cells, likely due to lower baseline phagocytic function and greater phenotypic deficiencies in mature cultures.

### Combined Autophagy Activation and PRPF31 Gene Augmentation Does Not Enhance Cellular Rescue

Building on our previous findings that autophagy activation via Rapamycin eliminates cytoplasmic aggregates and improves RP11-RPE cell survival (5), we investigated whether a synergistic approach, combining Rapamycin with *PRPF31* gene augmentation, could further enhance cellular rescue. To evaluate this, mature RP11-RPE cells were treated with PRPF31.AAV for two weeks, with the addition of Rapamycin during the final week of culture, followed by immunofluorescence analysis (**Figure S4**).

Consistent with our earlier results, PRPF31.AAV significantly restored nuclear PRPF31 localisation and nuclear speckle formation, reduced cytoplasmic aggregates, and increased cilia length and incidence compared to vehicle (DMSO) treated control. However, the addition of Rapamycin provided no measurable improvement beyond the effects achieved by PRPF31.AAV alone (**Figure S4**). While Rapamycin independently reduced RLBP1 and HSPB1 aggregates, and increased cilia length and its incidence, its combination with gene augmentation did not yield a cumulative reduction. Collectively, these data indicate that PRPF31 gene augmentation is the primary driver of cellular rescue in mature RP11-RPE cells, with no significant additive benefit from exogenous autophagy activation. Furthermore, this suggests that the therapeutic effects of PRPF31 restoration may already encompass the pathways activated by Rapamycin.

## Discussion

Mutations in core spliceosomal components, including the *PRPF* gene family, are well-established causes of autosomal dominant RP (5). Specifically, *PRPF31* mutations lead to the RP11 phenotype through a mechanism of haploinsufficiency, wherein the remaining wild-type expression levels are inadequate to sustain normal spliceosomal function. This impairment triggers a cascade of downstream cellular and functional defects, as previously characterised in RP11-iPSC-derived RPE cells and photoreceptors (11, 20). Consequently, AAV-mediated gene augmentation to restore *PRPF31* levels represents a highly rational therapeutic strategy, designed to recover splicing activity and reverse the RP11 disease phenotype.

In this study, we evaluated the therapeutic efficacy of AAV-mediated *PRPF31* gene augmentation in patient-specific iPSC-derived RPE and RO models, integrating transcriptomic, proteomic, and functional analyses to elucidate the mechanistic basis of rescue. We demonstrated that AAV delivery not only drives a robust increase in *PRPF31* mRNA and protein expression but also restores endogenous patterns of nuclear localisation, required for its function in the spliceosome. This molecular recovery was evidenced by normalised SC35 nuclear speckles architecture, more abundant and prominent foci of active spliceosomes in treated ROs, marked by p-SF3B1, and preponderance of pre-mRNA splicing pathway amongst the differentially spliced genes, collectively signalling a rescue of spliceosomal activity. The functional consequences of this rescue were evident at the transcriptome level: differential exon usage analysis identified over 500 alternative splicing events in both RPE cells and ROs following PRPF31 augmentation. In treated RPE cells, differentially spliced transcripts were enriched for processes central to mRNA splicing and processing, suggesting a positive feedback loop that reinforces homeostatic splicing patterns. Among these transcripts was *RBM5*, an important regulator of alternative splicing, which was affected at both the splicing level and in protein abundance. RBM5 is best known for its role in the splicing of pro-apoptotic genes (26). In treated ROs, differentially spliced transcripts were strikingly associated with the photoreceptor cilium, mitochondria, and inner segment - the compartments where protein trafficking, metabolic activity, and energy production are most vital. This compartment-specific enrichment provides a mechanistic link between corrected splicing and the structural integrity of photoreceptors. Furthermore, the downstream consequences of restored splicing fidelity were further revealed by integrated transcriptomic and proteomic profiling. Crucially, PRPF31 gene augmentation led to clearance of cytotoxic cytoplasmic aggregates across both RPE and RO models. This finding was corroborated by transcriptomic data showing the upregulation of amyloid-beta clearance and lipoprotein receptor-mediated endocytosis pathways. These transcriptional shifts aligned with proteomic evidence of the coordinated upregulation of clathrin-mediated endocytosis and lysosome components (e.g. CLTA, CLTB, NPC2, G2MA) and autophagy regulators (CALM3, CHMP2B, VPS37A). Notably, an essential regulator of POS phagocytosis by RPE cells, CDC42 was also upregulated at the protein level (22, 32). Ultimately, these molecular shifts translated into a multifaceted functional rescue. This was characterised by enhanced apical–basal polarity, elongated primary cilia, and re-established phagocytic capacity in RPE cells, alongside significantly increased light-evoked activity in photoreceptors, findings that align with the enrichment of synaptogenesis and ciliary pathways observed in our transcriptomic data. A recent study by Kaur et al., demonstrated that photoreceptors can reverse apoptotic features following stress removal, with mitophagy playing a central role in this recovery (33). This mechanistic finding aligns with our transcriptomics and proteomics data in PRPF31.AAV-treated ROs, which show regulation of genes critical for mitochondrial function and autophagy, potentially by correcting splicing of their transcripts. The upregulation of RRM2B and FDXR at the protein level is particularly notable, as deficiencies in these proteins cause mtDNA depletion or mitochondrial dysfunction, both of which lead to photoreceptor degeneration (27, 28).

Recently, our team reported the activation of autophagy via rapamycin to enhance the degradation of misfolded proteins in RP11-RPE cells (11). However, while rapamycin reduced the burden of cytoplasmic aggregates, its combination with AAV-*PRPF31* did not yield a significantly greater rescue effect than gene therapy alone. This suggests that while autophagy activation may alleviate aspects of cellular stress, it does not address the upstream driver of the disease as effectively as direct *PRPF31* restoration. Furthermore, PRPF31 restoration itself may normalise autophagic flux, potentially by correcting the splicing of autophagy-related genes identified in our proteomic data to be upregulated after gene therapy.

Our findings align with and extend recent studies highlighting the therapeutic potential of *PRPF31* gene augmentation across various RP11 models. Previous *in vitro* work using AAV2/Anc80-*PRPF31* successfully rescued phagocytic capacity and ciliogenesis in patient-derived RPE cells (8). Similarly, AAV2-7m8-*PRPF31* has been shown to enhance photoreceptor survival and arrest degeneration in ROs (17). These *in vitro* successes are further supported by *in vivo* data, where subretinal delivery of AAV2-7m8-*PRPF31* in *Prpf31* knockout mice preserved retinal structure and prevented pathological pigmentation (18). While AAV2/Anc80 and AAV2-7m8 were specifically designed to optimise retinal transduction, particularly in photoreceptors (34, 35), we sought to further refine this approach to target both RPE cells and photoreceptors. In this study, we employed the ShH10(Y445F) serotype, an AAV6-derived variant known for its superior efficiency in dual-targeting both iPSC-derived photoreceptors and RPE cells (36). This choice allowed for a more comprehensive rescue across the primary cell types affected in RP11.

While previous *in vitro* studies have largely assessed gene therapy efficacy in isolation, focusing either on RPE cells or photoreceptors (8, 17) our study utilised an integrated approach. By employing both RP11 iPSC-derived RPE and photoreceptor-containing ROs, we provide a more translationally relevant evaluation of therapeutic success. Furthermore, we conducted a systems-level investigation to gain deeper mechanistic insight into the rescue mediated by PRPF31.AAV gene therapy. Beyond confirming *PRPF31* overexpression and the rescue of phagocytosis and ciliogenesis, we characterised key subcellular hallmarks of RP11 that remained unaddressed in prior gene therapy reports. These include the restoration of nuclear *PRPF31* localisation and speckle organisation and the clearance of cytoplasmic aggregates. Ultimately, the systems-level validation provided by our integrated transcriptomic and proteomic analyses demonstrates that PRPF31 augmentation fundamentally reshapes the cellular landscape, upregulating pathways essential for energy metabolism, protein and organelle quality control, and visual function while suppressing inflammatory and neurodegenerative signatures. RP11 arises from the loss of a single functional *PRPF31* allele, leading to haploinsufficiency rather than the expression of a dominant-negative protein. This dosage-dependent mechanism is further evidenced by the incomplete penetrance observed in asymptomatic carriers, where higher expression of the remaining wild-type allele appears sufficient to preserve retinal function (10, 37). Our findings provide compelling evidence that therapeutically augmenting *PRPF31* expression recapitulates this protective effect, driving molecular, cellular, and functional recovery in both RPE and ROs.

Therapeutic timing remains a critical consideration for clinical translation. Late-stage PRPF31-RP is characterised by irreversible photoreceptor loss, a point beyond which gene augmentation cannot recover lost cells (38). We therefore evaluated whether *PRPF31* augmentation could rescue phenotypes in mature RPE cells, a model that recapitulates ageing-associated features, including accumulation of intracellular debris. Despite lower transduction efficiency in mature cells compared to earlier developmental stages, substantial phenotypic improvements were still achieved. This indicates that gene therapy may ameliorate defects even in later disease stages; preserving RPE structure and function, even partially, could be sufficient to delay further degeneration (8). This aligns with findings in *RPGR* X-linked RP models, where gene therapy slowed degeneration at intermediate or late stages, provided viable photoreceptors remained (39).

Together, these observations emphasise the importance of early intervention while also highlighting the potential for therapeutic benefit in more advanced cases. Given that patients typically seek treatment after symptom onset, late-stage intervention would focus on preserving remaining vision and slowing disease progression (40). Further investigation is required to define the optimal therapeutic window for maximal functional rescue in RP11.

In summary, our study provides a comprehensive, mechanistic evaluation of AAV-mediated *PRPF31* gene augmentation in human iPSC-derived RPE cells and photoreceptors. We demonstrate that increasing PRPF31 levels corrects the primary splicing defect, which in turn reshapes the transcriptomic and proteomic landscape to reverse the downstream cellular pathologies of RP11. The efficacy observed even in late-stage models suggests potential benefit across disease stages. Future *in vivo* studies should focus on optimising dosage, delivery timing, and long-term safety to translate these findings toward clinical application. Additionally, the mechanistic insights gained here, particularly the coordinate regulation of autophagy, endocytosis, and mitochondrial function downstream of splicing correction, may inform the development of combinatorial therapies for patients with advanced disease or those ineligible for gene therapy.

## Methodology

### Human iPSCs

The iPSC samples used in this study were obtained with informed consent according to protocols approved by Yorkshire and the Humber Research Ethics Committee (REC ref. no. 03/362) (20). In this study, we used PRPF31-iPSCs derived from three patients with severe (RP11S3, RP11S1) and very severe (RP11VS) phenotypes as well as unaffected controls (WT1, WT3) as described in earlier work (20). RP11S1 and RP11VS harbour the same *PRPF31* mutation (c.1115_1125 del11), whereas RP11S3 patients carry a different mutation (c.522_527+10del).

### iPSC culture

Human iPSCs were cultured on growth factor-reduced Matrigel (Corning, 354230) coated six-well plates in mTeSR1 media (StemCell Technologies, 85872) supplemented with 1% penicillin/streptomycin. Passaging was carried out using Versene (EDTA 0.02%) (Gibco, 15040-033) solution for 3-5 minutes. All cultures were maintained at 37 °C, in a humidified environment with 5% CO_2_. iPSCs were cryopreserved using freezing media containing 90% foetal bovine serum (Gibco, A5256801), 10% dimethyl sulfoxide (DMSO) (Sigma, D2650) and 10 µM ROCK inhibitor (Y-27632, Chemdea, CD0141).

### Differentiation of iPSCs to ROs

Generation of ROs derived from iPSCs was based on a previously described protocol (41). Briefly, iPSCs were dissociated to single cells using Accutase (Gibco, A1110501) and transferred to lipidure-coated 96-well U-bottomed plates at a density of 7000 cells/well in mTeSR1 media. At 48 hours, differentiation media was added to the cells, consisting of 41% IMDM (Gibco, 12440–053), 41% HAM’s F12 (Gibco, 31765–029), 15% Knockout Serum (KOS) (Gibco, 10828–028), 1% GlutaMAX (Gibco, 35050–038), 1% Chemically defined lipid concentrate (Thermo, 11905031), 1% Penicillin/Streptomycin (Gibco, 15140–122), 225 µM 1-Thioglycerol (Sigma, M6145). This is referred to as day 0 of differentiation. On day 6, the differentiation media was supplemented with 1.5 nM BMP4 (R&D, 314-BP). From day 18, the culture media was changed to DMEM/F12 (Gibco, 31330–038) supplemented with 10% fetal bovine serum (Gibco, A5256701), 1% N2 supplement (Gibco, 17502001), 0.1 mM Taurine (Sigma, T8691), 0.25 µg/ml Fungizone (Gibco, 15290018), 1% Penicillin/Streptomycin (Gibco, 15140–122) and 0.5 µM Retinoic Acid (Sigma, R2625) until day 120.

### Differentiation of iPSCs to RPE cells

iPSCs were differentiated to RPE cells by following a directed differentiation protocol as described in a previous publication (42). Briefly, once the iPSC colonies reached 80-90% confluency, mTeSR1 media was replaced with differentiation medium (DMEM/F12 + GlutaMAX (Gibco, 31331-028), supplemented with 50 µM 2-Mercaptoethanol (Gibco, 21985023), 1% MEM NEAA (Gibco, 11140035), 20% KOS (Gibco, 10828–028), 10mM Nicotinamide and 1% penicillin/streptomycin). At day 7, nicotinamide was substituted with 100 ng/mL Activin A (PeproTech, 120-14E) for 7 more days. Thereafter, 3 µM CHIR99021 (Sigma, SML1046) replaced Activin A until day 42. Cells were cultured in media consisting of DMEM/F12 + GlutaMAX, 50 μM 2-Mercaptoethanol, 1% MEM NEAA, 4% KOS and 1% Penicillin/Streptomycin until harvesting the RPE patches around day 90. The collected patches were dissociated in TrypLE Enzyme (10X) (Gibco, A12177-01) for 20 minutes at 37°C and replated at a density of 1.5 x 10^5^ cells per cm^2^ on 12-well plates or on 24-Transwell inserts (Greiner Bio-one, 662641).

### Production of recombinant AAV viral vector

Adeno-associated viral vectors were packaged in serotype ShH10(Y445F) (Y445F) capsid (43). The transgenic CMV.PRPF31 construct was prepared by cloning the *PRPF31* coding sequence into the multiple cloning site of pD10 backbone, upstream of an SV40 polyadenylation signal. An equivalent CMV.EGFP construct was used as a negative control, and sequences were confirmed by nanopore sequencing (FullCircle laboratories). AAV particles were prepared using a triple-transfection protocol using HEK293T cells AAVPro (TakaraBio), and were isolated, purified on AVB columns and quantified by qPCR as previously described (44).

### AAV transduction in RPE cells

To mimic the “early” stage AAV transductions, iPSC-RPE cells were passaged on 24-Transwell inserts 3 days before transduction, while for “late” AAV transduction, iPSC-RPE cells were matured on transwell inserts for more than 12 weeks before AAV supplementation. ShH10(Y445F).CMV.GFP or ShH10(Y445F).CMV.PRPF31 AAVs were suspended in RPE media to achieve an MOI of 100,000 vector genomes (vg)/cells in a final volume of 50 µl. The AAVs were added to the RPE transwells apically and incubated overnight at 37°C with 5% CO_2_. Media was added to the transwells the next day and changed twice a week afterwards. All transduced RPE cells were cultured for two weeks before collection for further assays.

### AAV transduction in ROs

ROs were cultured to day 180 before transduction with ShH10(Y445F).CMV.GFP.AAVs or ShH10(Y445F).CMV.PRPF31.AAVs at the MOI of 3×10^6^ vg/cell in a final volume of 50 µl. ROs were incubated overnight with AAVs and subsequently cultured for two weeks.

### RNA extraction and reverse transcription

Samples were washed with Phosphate-Buffered Saline (PBS) before lysis with RNA Lysis buffer from the RNA extraction kit ReliaPrep RNA Cell Miniprep System (Promega, Z6010). Total RNA was extracted as per the manufacturer’s instructions, including a DNase incubation step to remove any contaminating DNA. Concentration of RNA was measured with a NanoDrop 2000 Spectophotometer (Thermo Scientific). Next, 1 µg of RNA was reverse transcribed into cDNA using random primers from the GoScriptTM Reverse Transcription System (Promega, A5000) following the manufacturer’s instructions.

### Quantitative real-time polymerase chain reaction

RT-PCR was performed using the GoTaq qPCR Master Mix (Promega, A6001) according to manufacturer’s instructions. C_t_ values were measured using the QuantStudio™ 7 Flex Real-Time PCR System (ThermoFisher) in a 10 µl reaction volume, with each sample run in triplicate. Relative gene expression to *GAPDH* was normalised to the control sample using the ΔΔCT method. All primer sequences are shown in **Table S5**.

### Western blotting

RPE cells and ROs were washed with PBS and re-suspended in RIPA lysis buffer (EMD Millipore, 20-188) supplemented with EDTA-free protease inhibitor cocktail (Roche, 04693159001). Total protein concentration of lysates was determined using the BCA Protein assay kit (Thermo Scientific, 23225). Samples were reduced and denatured using NuPAGE LDS Sample Buffer (4x) (Invitrogen, NP0007) and NuPAGE Sample Reducing Agent (10x) (Invitrogen, NP0004), followed by a 10-minute incubation at 70°C. Ten µg of each lysate were then applied to sodium dodecyl sulfate-polyacrylamide gel electrophoresis (SDS-PAGE) (Invitrogen™, NW04120BOX). Afterwards, the gels were transferred to Hybond polyvinylidene difluoride (PVDF) membranes (Invitrogen, IB24002) and blocked for 1 hour in 5% dried skimmed milk in TBST (1% Tween 20, 5% tris-buffered saline (TBS) in dH_2_O). Membranes were immunostained using antibodies against PRPF31, GFP and GAPDH (**Table S5**). Horseradish peroxidase (HRP)-conjugated secondary antibodies were used to detect bound primary antibodies (**Table S5**). The signal was detected using the SuperSignal West Pico PLUS Chemiluminescent Substrate (ThermoFisher Scientific, 34580) and was visualised by Amersham Imager 600 imager. Quantification of the signal was carried out with FIJI software (45).

### Immunofluorescence analysis in RPE cells

RPE cells were fixed with 4% PFA (paraformaldehyde) (Sigma, 47608) for 30 minutes at room temperature or with 5% methanol for 5 minutes at -20 °C for ARL13B staining. PFA was removed and samples were rinsed with PBS twice and stored at 4 °C until required. Samples were blocked in PBS with 10% serum and 0.3% Triton-X-100 (Sigma, T8787) for 1 hour at room temperature before incubation with primary antibodies at 4 °C overnight **(Table S5)**. Following three PBS washes, samples were incubated for 1 hour with secondary antibodies in PBS and 1% serum and later with DAPI stain for 20 minutes. For ARL13B staining, 1% marvel in PBS was used for blocking samples and antibody incubation. RPE cell samples were then mounted with Vectashield and imaged using the Axio Imager 4 upright microscope with Apotome structured illumination fluorescence using 20x and 63x oil objectives (Zeiss, Germany) obtaining 5-10 images per sample, with z-stack acquisitions of 20-30 optical sections per image (0.49 µm interval). Images were processed in ZenBlue software (Zeiss, v.2.5) and presented as a maximum intensity projection.

### Immunofluorescence analysis in ROs

ROs were fixed with 4% PFA for 15 minutes at room temperature. Following PBS washes, ROs were incubated at 4°C overnight in 30% sucrose in PBS, embedded in OCT Embedding matrix (CellPath, KMA-0100-00A) and cryosectioned at 10 µm thickness. Sections were incubated with blocking solution (10% serum, 0.3% Triton-X-100) for 1 hour at room temperature, followed by overnight incubation at 4°C with primary antibodies (**Table S5**) in antibody diluent (1% serum, 0.1% Triton-X-100 in PBS). After PBS washes, sections were incubated with secondary antibodies (**Table S5**) conjugated to fluorophores for 1 hour. Nuclei were labelled with DAPI nuclear stain for 20 minutes. Sections of ROs were sealed with Vectashield and imaged using the Axio Imager-4 upright microscope with Apotome structured illumination fluorescence. For each sample, 5 organoids were included, with 5-12 images acquired per sample and with z-stack acquisitions of 20-25 optical sections per image. ZenBlue software (Zeiss, v.2.5) was used to process images, and they are presented as a maximum intensity projection.

### Phagocytosis assay in RPE cells

Bovine rod photoreceptor outer segments (POSs) were labelled with IncuCyte pHRodo red cell labelling dye (Sartorius, 4649) for 1 hour at room temperature in the dark with shaking, as per manufacturer’s protocol. Labelled POSs were then washed with PBS and resuspended in RPE media supplemented with 1% fetal bovine serum. 20 POSs/cell were added to the RPE cells in a total volume of 75 µl at 17 °C for 30 minutes followed by incubation in IncuCyte Live imaging system. Images were taken every 30 minutes for 4 hours total. Quantification was carried out using the orange image mean intensity and the percentage of red area at 4 hours, calculated using ImageJ (45).

### Multielectrode array recordings in ROs

Multielectrode array (MEA) recordings were performed on the BioCamX platform using the high-density MEA CorePlate™ 1W 27/42L (3Brain GmbH, Lanquart, Switzerland), which integrates 4096 square microelectrodes in a 2.67 × 2.67 mm area and 42 μM inter-microelectrode spacing. 24 hours prior to MEA recordings, 9-cis retinal (10 nM; Sigma-Aldrich, UK) was added to the incubation medium. Organoids were transferred to 33 °C artificial cerebrospinal fluid (aCSF) containing the following (in mM): 118 NaCl, 25 NaHCO_3_, 1 NaH_2_PO_4_, 3 KCl, 1 MgCl_2_, 2 CaCl_2_, 10 glucose, 0.5 L-Glutamine, and 0.01 9-cis-retinal. Organoids were opened longitudinally and placed with the presumed retinal ganglion cell (RGC) layer facing down onto a 4096-channel multielectrode array (MEA 3Brain GmbH, Lanquart, Switzerland). They were flattened using a translucent polyester membrane filter (Sterlitech Corp., Kent, WA, USA) with a stainless-steel ring of 3 mm diameter on top. The organoids were allowed to settle for at least 2 hours while continuously perfused (1 mL/min) with bubbled aCSF. Recordings are filtered with a 50 Hz high-pass filter in BrainWaveX (3Brain) and stored in an HDF5 format. Action potentials (spikes) from retinal organoid ganglion cells were detected and sorted using Herdingspikes2 (https://github.com/mhhennig/HS2) as previously described (46). Briefly, spikes were first identified as threshold crossings on each channel, then merged into unique events based on spatial and temporal proximity. For each detected spike, a location was estimated from the signal’s center of mass. Spike sorting involved clustering all events using a feature vector that includes the locations and the first two principal components of the largest waveform.

Light stimuli were projected onto the retina as described elsewhere (47). Briefly, light stimuli were projected using a DLP LightCrafter (Texas Instruments, TX, USA) and custom optics onto the 2.67 x 2.67 mm field of the MEA chip, where the retinal organoids are placed. A full field stimulus with increasing contrasts (1-second contrast steps, max irradiance 217 μW/cm2, 1 second black, followed by 1 second 33% grey, 1 second black, 1 second 66% grey, 1 second black, and 1 second white) was used to detect potential basic light-driven responses in the retinal organoids. cGMP (cyclic guanosine monophosphate) acts as a second messenger in photoreceptor cells, regulating the phototransduction cascade. Statistical significance and firing rate analyses were evaluated by using MATLAB (Mathworks, MA, USA) and Prism (GraphPad, CA, USA). RGCs were considered responsive if they showed at least a 25% increase or decrease in spiking activity during a 90 s time window during stimulus (light onset) compared to a similar time window before the stimulus (spontaneous activity). For each cell, all spikes occurring during these time windows were counted, and the median percentage change (± IQR) in activity between windows was calculated.

### RNA-Seq Data analyses

#### Quality control and alignment against the reference genome

RNA paired-end sequencing quality control was assessed through FastQC (www.bioinformatics.babraham.ac.uk/projects/fastqc) and multiQC (48). An average of 125 million reads were sequenced per sample. All libraries were mapped against the Human GRCh38 release-109 reference genome (retrieved from http://ftp.ensembl.org/pub/release-109/fasta/homo_sapiens/dna/Homo_sapiens.GRCh38.dna.primary_assembly.fa.gz) through STAR aligner (49) with default parameters. STAR-generated sorted BAM output files were used for assigning read counts to gene features with featureCounts (50) with the following parameters: -p -C -M -O --fraction -s 2. We relied on Homo_sapiens.GRCh38.109.gtf annotation file downloaded from the same ensembl ftp address above.

#### Differential gene expression

Read counts’ table generated by featureCounts was then used as input for differential expression (DE) analyses relying on the DeSeq2 negative binomial distribution model (51) through a local fitting type and a 0.05 FDR threshold. DE pairwise comparisons were performed, always setting PRPF31/GFP as the fold-change ratio to be assessed. *varianceStabilizingTransformation(dds, blind=T)* and *plotPCA()* functions from the DESeq2 package were run on the normalised counts matrix for Principal Component Analyses. EnhancedVolcano (https://bioconductor.org/packages/release/bioc/html/EnhancedVolcano.html) was employed for an overall DE visualisation through volcano plots. All tools described in this paragraph were run under the R environment version 4.4.

#### Differential splicing

STAR-generated sorted BAM files were used as input for rMATs turbo v4.1.2 (52) for differential splicing (DS) analyses under the following parameters: --libType fr-firststrand --readLength 150 --variable-read-length –novelSS. Sample groups’ pairwise comparisons for DS analyses followed the same rationale as for DE ones described above. An *ad-hoc* PERL script (https://github.com/eltonjrv/bioinfo.scripts/blob/master/filter_rMATs.pl) was written for filtering rMATs results according to our set of pre-established thresholds: FDR < 0.05, Inclusion Level Difference (IncLevelDifference) > 0.05, and both inclusion and skipping junction counts average (IJC_SJC_avg) > 5.

#### Gene enrichment functional analysis

GO-, KEGG-, and Reactome-based gene enrichment analyses for individual lists of either DE or DS genes were performed by clusterProfiler Bioconductor package (53) under R v4.4, setting adjusted p-value < 0.05 on *enrichGO(), enrichKEGG(), and enrichPathway()* functions, respectively.

Table of software according to Cell Press Key Resources:

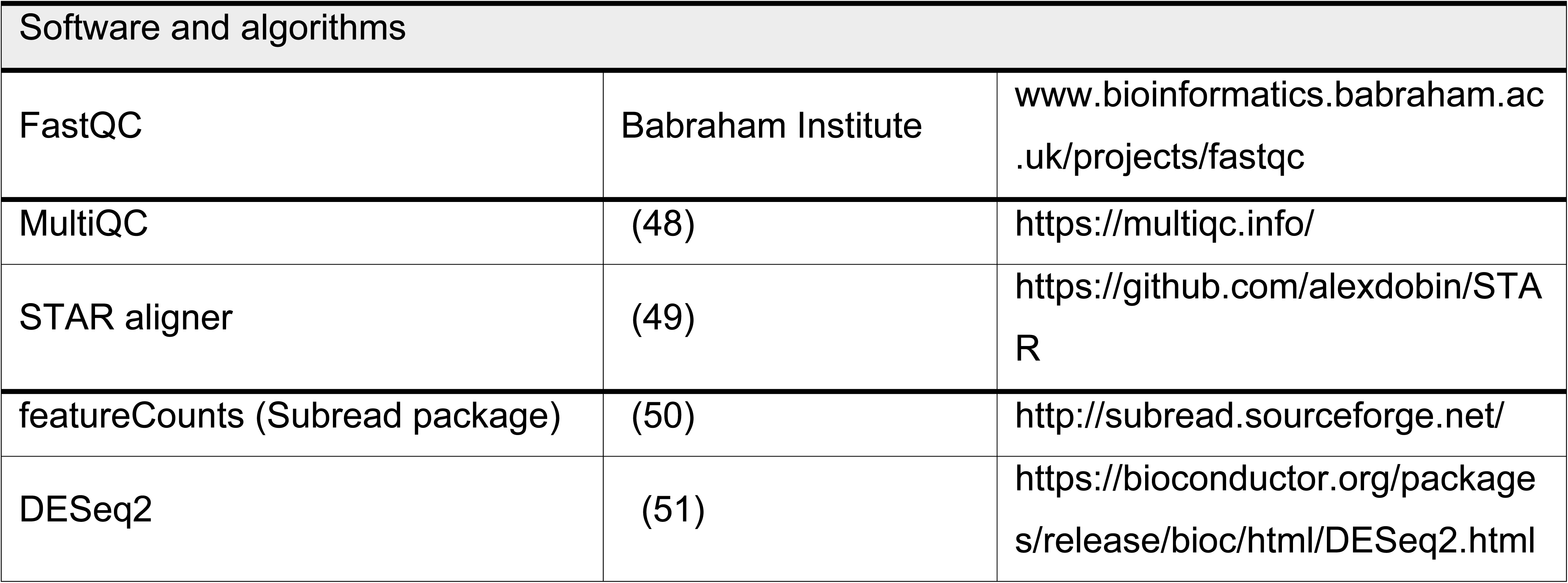
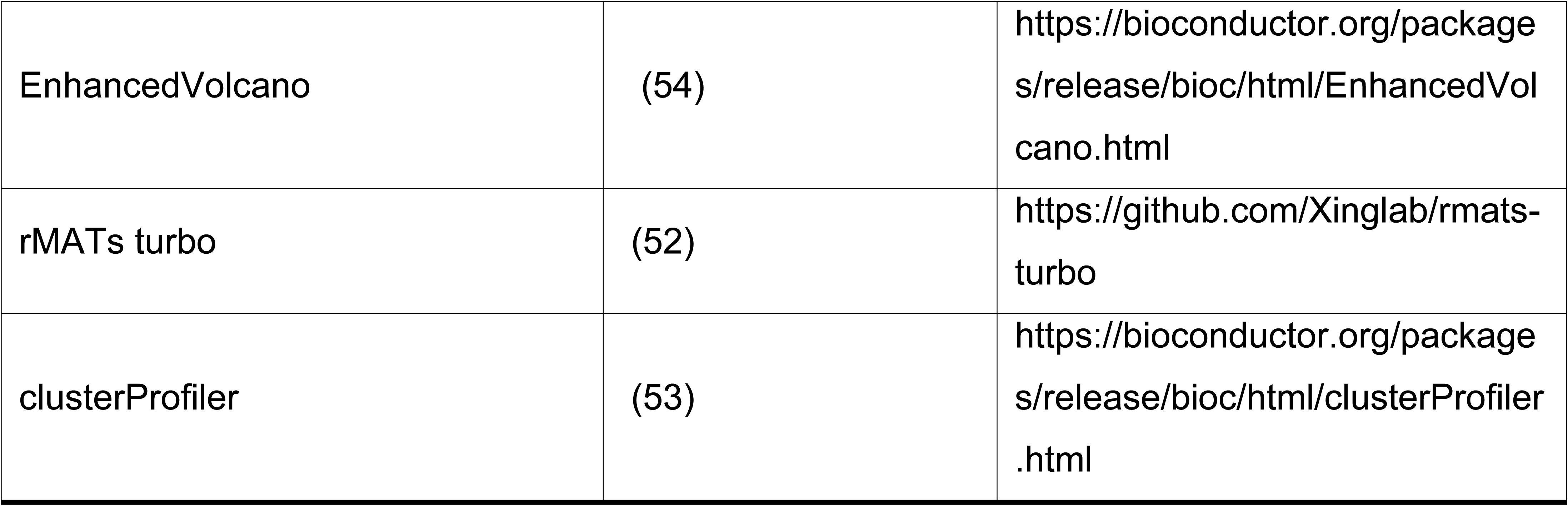

| Software and algorithms |  |  |
| --- | --- | --- |
| FastQC | Babraham Institute | www.bioinformatics.babraham.ac.uk/projects/fastqc |
| MultiQC | (48) | https://multiqc.info/ |
| STAR aligner | (49) | https://github.com/alexdobin/STAR |
| featureCounts (Subread package) | (50) | http://subread.sourceforge.net/ |
| DESeq2 | (51) | https://bioconductor.org/packages/release/bioc/html/DESeq2.html |
| EnhancedVolcano | (54) | https://bioconductor.org/packages/release/bioc/html/EnhancedVolcano.html |
| rMATs turbo | (52) | https://github.com/Xinglab/rmats-turbo |
| clusterProfiler | (53) | https://bioconductor.org/packages/release/bioc/html/clusterProfiler.html |

### Proteomics analysis

#### TMT labelling for mass spectrometry

RPE cells or ROs were lysed by boiling in 100 µL 2% (w/v) SDS, 0.5 mM EDTA, 50 mM HEPES, pH 8.0, 1x HALT phosphatase and protease inhibitor cocktail (Thermo Scientific) for 5 minutes at 95°C. Lysates were sonicated in a Bioruptor device (Diagenode) for 5 minutes with 30 seconds on-and-off cycles, centrifuged for 10 minutes at 15,000 g and clear supernatants were transferred to new sample tubes. Protein concentrations were determined using the Pierce BCA protein assay kit (Thermo Scientific), and 100 µg of total protein samples were further processed for isobaric tandem mass tag (TMT) labelling. Briefly, proteins were reduced with 10 mM Tris-(2-carboxymethyl)-phosphine hydrochloride (TCEP) and alkylated with 40 mM 2-Chloroacetamide (CAA) for 30 minutes at 37°C followed by SP3 clean-up using Sera-Mag Carboxylate SpeedBeads (E3 & E7; Cytiva). On-bead tryptic digestion (1:20 enzyme-to-protein ratio) in 100 mM tetraethylammonium boride (TEAB) buffer was performed overnight at 37°C. Protein digests were split into 25 µg aliquots for TMT labelling and dried by vacuum centrifugation in a SpeedVac (Thermo Fisher Scientific). Additionally, 5 µg aliquots of each sample (separately for RPE cells and retinal organoids) were combined to use as a reference channel for TMT labelling. The reference sample was split into 25 µg aliquots before drying in a SpeedVac.

Samples were labelled using the TMTpro 16plex Label Reagent Set (Thermo Scientific; 0.5 mg vials) according to the manufacturer’s instructions except for using a sample-to-tag ratio of 1:4 (25 µg peptides labelled with 100 µg TMT). Due to the large number of samples, they were split into two sets and separately labelled with TMTpro reagents. The reference sample was labelled with TMTpro-134N in both sets of samples.

Following the one-hour incubation at room temperature, labelling reactions were quenched by 5% hydroxylamine for 15 minutes. Labelled peptide samples of each TMT set were combined and dried in a SpeedVac. Samples were then re-dissolved in 4% (v/v) acetonitrile (ACN), 0.1 % (v/v) trifluoroacetic acid (TFA) in water and cleaned up using C18 Micro Spin Columns (Harvard Apparatus). To remove unbound TMT labels, two washing steps with 4% (v/v) ACN, 0.1 % (v/v) TFA in water were performed following protein binding. TMT-labelled peptides were eluted with 150 µL 50% (v/v) ACN, 0.1% (v/v) TFA in water, followed by 150 µL 80% (v/v) ACN, 0.1 % (v/v) TFA in water. Next, peptides were dried by vacuum concentration in a SpeedVac.

To reduce sample complexity and increase peptide detection during mass spectrometry (MS) measurement, TMT-labelled peptide samples were fractionated by high-pH reversed-phase chromatography. Briefly, samples corresponding to ∼ 75 µg labelled peptides dissolved in 30 µL of 2% (v/v) ACN, 10 mM ammonium hydroxide (NH_4_OH) were injected into an X-Bridge® C18 column (3.5 μm, 1.0 x 150 mm; Waters) and separated using a step gradient of 8 – 95% (v/v) buffer B (80% (v/v) ACN, 10 mM NH_4_OH in water) over 50 min. One-minute fractions were collected and combined to 20 fractions by concatenating. Fractions were then dried in a SpeedVac and re-dissolved in 20 µl of 2% (v/v) ACN, 0.05 % (v/v) TFA in water for LC-MS/MS analysis.

#### LC-MS/MS analysis

Fractionated TMT samples were analysed by data-dependent acquisition (DDA) mass spectrometry using an Orbitrap Exploris 480 mass spectrometer (Thermo Scientific) coupled to an Ultimate 3000 RSLCnano UHPLC system (Thermo Scientific) in injection duplicates. Dried peptides were re-dissolved in 2% (v/v) ACN, 0.05% (v/v) TFA in water and injected into the UHPLC system, where they were concentrated on a C18 PepMap100-trapping column (0.3 x 5 mm, 5 μm; Thermo Fisher Scientific) and separated on an in-house packed analytical C18 column (75 μm x 300 mm; Reprosil-Pur 120C18-AQ, 1.9 μm; Dr. Maisch GmbH) using a 100 min linear, step gradient of 12-50% (v/v) buffer B (80% (v/v) ACN, 0.1% (v/v) formic acid (FA) in water) at 300 nL/minute (step 1: 12-36% buffer B over 87 minutes; step 2: 36%-50% buffer B over 13 minutes). Precursor ion scans were acquired at a resolution of 120,000 with an AGC target of 3E6 and maximum injection time of 60 ms in the orbitrap mass analyser across a scan range of 350-1,650 m/z. Precursor ions with a charge state of 2-7 were selected with an isolation window of 0.8 m/z and fragmented by higher-energy collisional dissociation (HCD) with a fixed collision energy of 32%. Fragment ion scans were acquired at a resolution of 60,000 with an AGC target of 2E5 and maximum injection time of 128 ms in the orbitrap mass analyser (turboTMT-mode off). The maximal cycle time was set to 3s, and dynamic exclusion was set to 20 s.

#### Data processing

MS/MS spectra were searched against the UniProtKB human reference proteome (release 2024-10-02; 20,668 entries) using MaxQuant version 2.6.5.0. The type “Reporter MS2” was selected and information about isotopic distributions for the TMTpro labelling reagents were included. Data for RPE cells and retinal organoids were searched separately. Experimental groups were defined as A and B representing the two different sets of TMT-labelled samples for RPE cells and retinal organoids, respectively. Fraction numbers were defined with different numbers for each experimental group. The searches were run with default settings with the following modifications: (i) Filtering by PIF was enabled with a minimum reporter PIF of 0.5; (ii) Trypsin was selected as digestive enzyme instead of Trypsin/P; (iii) Writing of mzTab tables was enabled; (iv) Match between runs was enabled for identifications.

Protein ratios were log2 transformed and median normalised using Perseus (v.2.0.6.0). The reported GFP.AAV/PRPF31.AAV ratios are the average of three biological replicates. To identify the differentially expressed proteins (DEP), those proteins with a mean log2 fold change (LFC) ≥ 0.26 or ≤ -0.26 were defined as DEP. Gene ontology (GO) enrichment analyses were carried out using DAVID and were visualised using SRplot.

### Quantification of PRPF31 cytoplasmic volume

The PRPF31 cytoplasmic volume was quantified from Z-stack images using Huygens Image Analysis Software. Around 9-14 images were analysed per sample. Briefly, the total volume and cytoplasmic volume were taken from each image using the object analyser to calculate the isosurface volume (IsoVol). The total volume was calculated by summing the individual volumes of selected objects within the specific channel of interest (PRPF31). Cytoplasmic volume was then calculated by subtracting the nuclear PRPF31 objects, which were co-localised with nuclei object. Nuclear volume was calculated by subtracting the cytoplasmic volume from the total volume. Nuclei were counted with ImageJ/FIJI and ratio of each volume were used to get percentage of nuclear volume per image (45).

### Quantification of nuclear speckles

The volume and intensity of nuclear speckles were quantified from Z-stack images using Huygens Image Analysis Software. The volume of SC35 was quantified by calculating the total isosurface volume, and the intensity by the sum of all voxel values for the signal. The results were normalised against the number of cells in each sample, counted with ImageJ/FIJI. The percentage of SC35-positive cells was quantified by ImageJ/FIJI and are presented as SC35-positive nuclei / total nuclei number % in each image (45). Around 7-9 images from each group were analysed.

### Quantification of aggregate volume

The total volume of aggregates (HSPB1, RLBP1) was quantified from Z-stack images, using Huygens Image Analysis Software. Analyses were performed on approximately 9–12 images per sample. HSPB1 and RLBP1 objects were segmented in Huygens by thresholding, and the object analyser was used to calculate the sum volume of all objects from each image. Nuclei were counted with ImageJ/FIJI and results are shown as the mean volume per cell in each area of interest (45).

### Quantification of cilia

The Z-stack images obtained were imported to Huygens essential software. ARL13B objects were segmented, and the length of objects was calculated with the object analyser. Nuclei were counted with Image J by thresholding and analysing the individual particles (DAPI)(45). The reported cilia length is the average length of all cilia in the image. The cilia incidence was calculated as cilia number / total cell number (DAPI) in the same region %. Quantification was performed on 7-9 images per sample.

### Rapamycin treatment

Both RPE cells and ROs were treated with 500 nM Rapamycin (AY-22989) (Selleck chemicals, S1039) suspended in media in a final volume of 150 µl for RPE cells and 50 µl for ROs three times a week for 1 week before sample collection. The final concentration of Rapamycin was prepared using dimethyl sulfoxide (DMSO).

### Statistical analysis

P-values for normally distributed data were calculated using one-way ANOVA with Dunnett’s post hoc test using GraphPad Prism Software Inc. (10.1.2) (San Diego, CA, USA). Data were plotted in GraphPad Prism, and error bars represent the standard deviation or as otherwise indicated. The statistical significance of pairwise comparisons shown on graphs is indicated by *p < 0.05, **p < 0.01, ***p < 0.001 and ****p < 0.0001.

## Data availability

RNA-Seq data were deposited to GEO, accession number GSE309908.

The mass spectrometry proteomics data have been deposited to the ProteomeXchange under the accession number PXD076430.

## Author contributions

Maria Elia - data curation, formal analysis, investigation, methodology, visualisation, writing – original draft, writing – review and editing.

Valda Pauzuolyte, Maria Georgiou, Mark Basche, Carina Hansohn, Robert Atkinson - data curation, formal analysis, investigation, methodology, writing – review and editing.

Maria Georgiou- data curation, formal analysis, investigation, methodology, writing – review and editing.

Elton Vasconcelos, Sushma Grellscheid - data curation, formal analysis, investigation, methodology, visualisation, writing – review and editing.

Colin A Johnson, Henning Urlaub - investigation, methodology, formal analysis, supervision, writing – review and editing.

Gerrit Hilgen - formal analysis, methodology, visualisation, writing – original draft, writing – review and editing.

Alexander Smith - formal analysis, methodology, visualisation, writing – review and editing. Sina Mozaffari-Jovin, Robin Ali, and Majlinda Lako - conceptualisation, formal analysis, funding acquisition, investigation, methodology, project administration, resources, visualisation, writing – original draft, writing – review and editing.

## Acknowledgements

We are grateful to Retina UK (GR # 601, #GR584, #GR595), Fight for Sight UK (1456/1457), MRC UK (MR/T017503/1), and EPSRC/ERC (EP/Y031016/1) for funding this work. The purchase of IncuCyte used in this study was supported by a UKRI MRC Capital Funding for World Class Labs Award (MR/X012360/1). HU was supported by the German Research Society (DFG) within the CRC SFB1289 (project number 317475864). We would also like to thank the Newcastle University FMS Bioimaging Unit, flow cytometry, and microscopy core facilities for their continued support during this project. We would like to thank LeedsOmics at the University of Leeds for help with technical assays and RNA-seq analysis, and University of Leeds alumnus funding from Peter & Sue Cheney.

