## Supplementary Information for "Cell-autonomous restoration of splicing homeostasis and RP11 phenotype in patient-derived RPE and retinal organoids by *PRPF31*.AAV gene therapy"

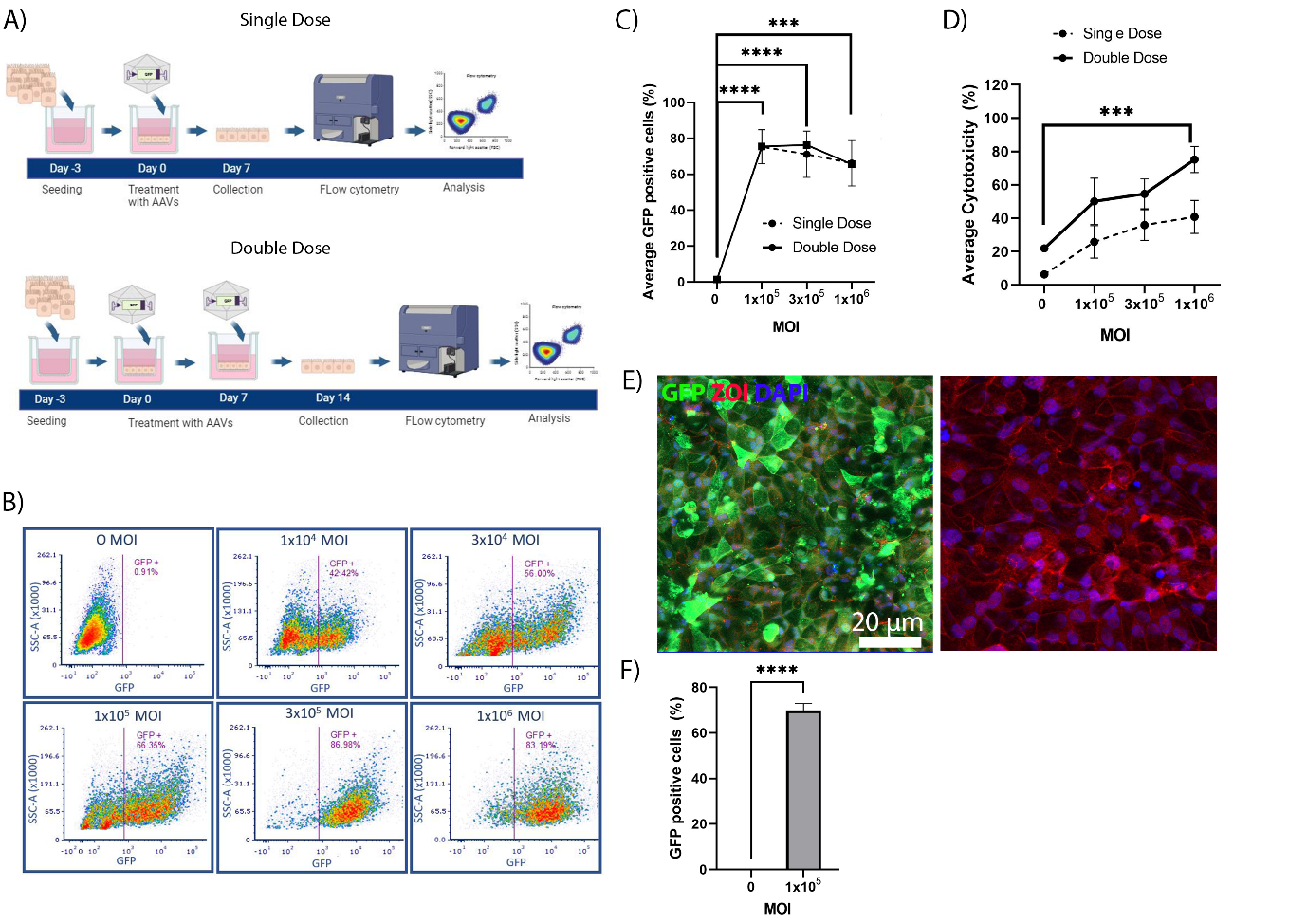
Supplementary information:

**Figure S1: Optimisation of ShH10.AAV Transduction Efficiency and Safety in iPSC-Derived RPE Cells.**

(A) Experimental workflow. Schematic representation of the transduction protocol for iPSC-derived RPE cells utilising single or double dosing regimens of ShH10.CMV.GFP.

(B) Single dose-response analysis via flow cytometry. Representative scatter plots and population gating illustrate the percentage of GFP-positive (GFP+) RPE cells across a MOI titration range of 0 to 1×10^6^ vg/cell.

(C) Transduction efficiency of single vs. double dosing. Comparative analysis of GFP+ RPE cells following a single dose (dotted line) or double dose (continuous line) of ShH10.CMV.GFP.

(D) Cytotoxicity profiling. Evaluation of cellular toxicity (%) across the three most efficient multiplicities of infection (MOIs) for both single and double dosing regimens. A significant increase in cytotoxicity was observed at the highest MOI in the double-dose group.

(E) Visualisation of viral transgene expression following single dosing with ShH10.CMV.GFP AAVs. Immunofluorescence microscopy confirms robust GFP expression (green) in RP11-RPE cells transduced with ShH10.CMV.GFP at an MOI of 1×10^5^ (left) compared to untreated controls (right). Nuclei are counterstained with DAPI (blue). Scale bar: 20 µm.

(F) Quantitative validation of GFP expression. Image-based quantification confirms a significant increase in GFP+ cells following treatment with MOI of 1×10^5^ vg/cell ShH10.CMV.GFP AAVs. Data are presented as mean ± SEM (n=3).

Statistical Analysis: Data in (C) and (D) were analysed using a two-way ANOVA followed by Šídák's multiple comparisons test (n=5). Statistical significance is indicated as follows: ***p<0.001, ****p<0.0001.

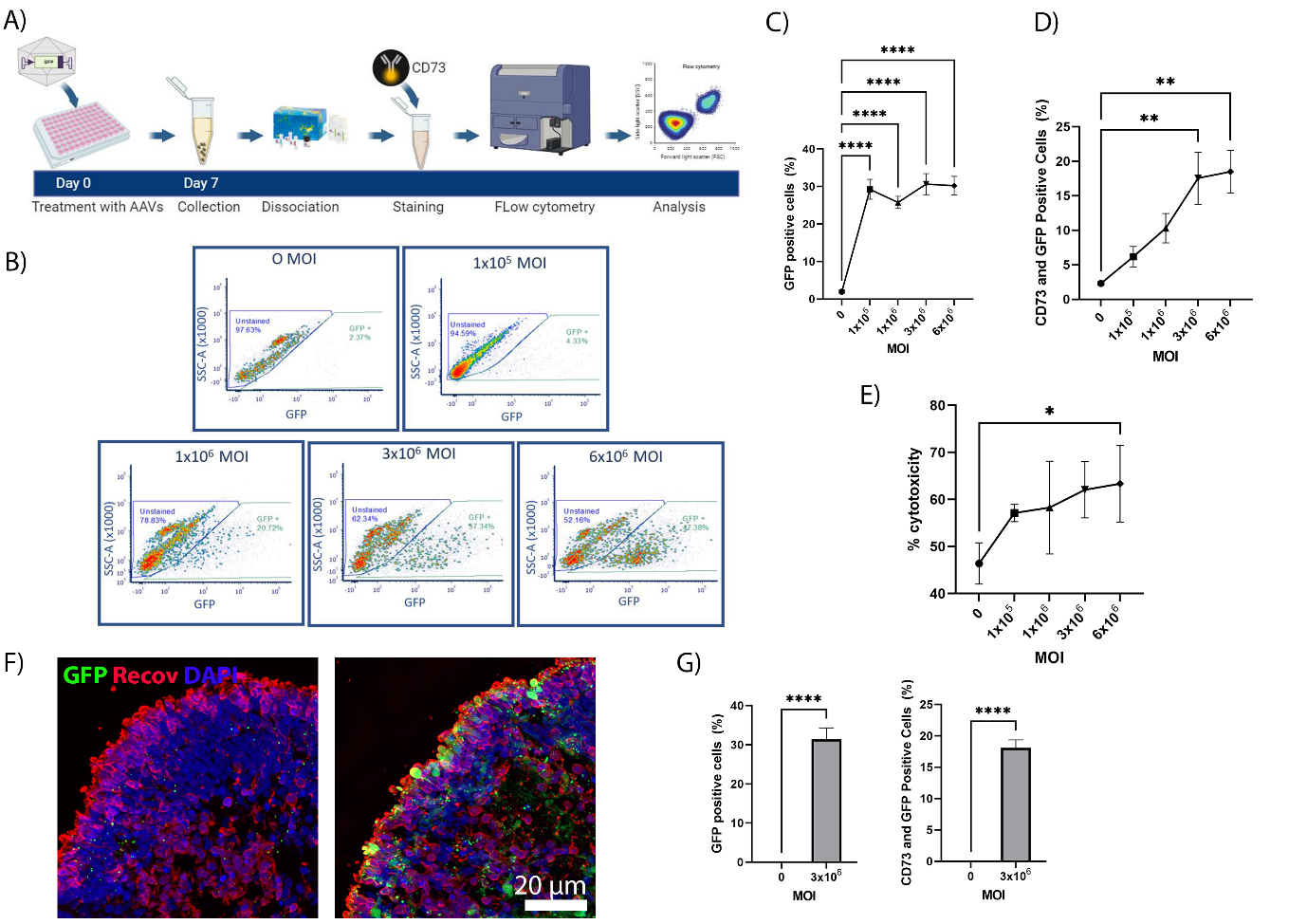

**Figure S2: Assessment of ShH10.AAV Transduction Efficiency and Photoreceptor Tropism in iPSC-derived ROs.**

(A) Experimental workflow for organoid transduction. Schematic of the protocol for treating day 150 (D150) iPSC-derived ROs with ShH10.CMV.GFP AAVs. To quantify efficiency, ROs were dissociated into single-cell suspensions and immunostained with the photoreceptor marker CD73 before flow cytometric analysis.

(B) Representative flow cytometry profiles. Gating strategy and fluorescence plots for ROs transduced with ShH10.CMV.GFP AAVs across a MOI range of 0-6x10^6^ vg/cell.

(C) Global transduction efficiency. Comparison of the percentage of GFP% cells in control and RP11-ROs across four MOIs. Statistical significance was determined via ordinary one-way ANOVA with Dunnett’s multiple comparisons test.

(D) Targeted photoreceptor transduction. Percentage of CD73+/GFP+ dual-positive cells, confirming ShH10 tropism for the photoreceptors residing in the apical aspect of ROs. Statistical significance was assessed by ordinary one-way ANOVA.

(E) Dosage-dependent cytotoxicity. Evaluation of LDH release/cytotoxicity (%) in ROs. Significant toxicity was observed only at the highest MOI (6x10^6^ vg/cell). Data were analysed using two-way ANOVA with Dunnett's multiple comparisons test (*p < 0.1).

(F) Histological validation of transgene expression. Immunofluorescence imaging of RO sections confirms high GFP expression (green) localised to the photoreceptor layer in transduced ROs (3x 10^6^ vg/cell, right panel) compared to untreated controls (left panel). Nuclei are counterstained with DAPI (blue). Scale bar: 20 µm.

(G) Quantitative image analysis. Volumetric quantification of GFP fluorescence confirms a significant increase in signal in ROs treated with 3 x 10^6^ vg/cell compared to controls. Data analysed via unpaired t-test.

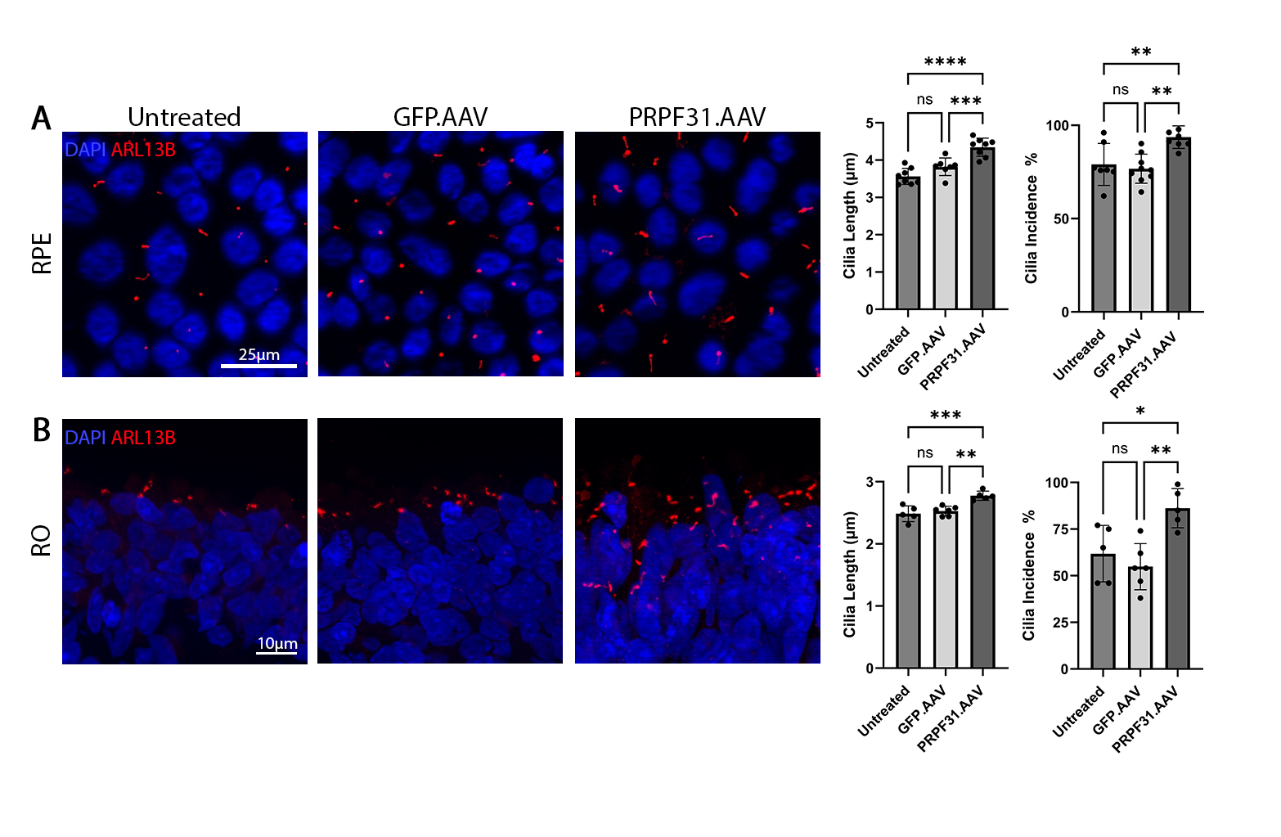
Statistical Analysis: All data are presented as mean ± SD, n = 3 (**p < 0.05, ****p < 0.0001).

**Figure S3: *PRPF31* Augmentation Rescues Cilia Incidence and Length in Both RP11-RPE Cells and ROs.**

(A) Ciliary rescue in RP11-RPE cells. Immunofluorescence imaging using the marker ARL13B (red) highlights the restoration of primary cilia in RP11-RPE cells following PRPF31.AAV treatment. PRPF31.AAV-treated RPE cells are characterised by increased cilia incidence (percentage of ciliated cells) and average cilia length compared to the untreated and GFP. AAV-treated RPE cells. Nuclei were counterstained with DAPI (blue). Scale bars: 25 µm.

(B) Recovery of cilia in RP11-ROs. Immunostaining of RO sections for ARL13B (red) demonstrates that PRPF31 augmentation reverses ciliary deficits in photoreceptors. Quantitatively, both cilia length and incidence are significantly increased compared to untreated and GFP.AAV-transduced controls. Nuclei were counterstained with DAPI (blue). Scale bars: 10 µm.

Statistical Analysis: Data are presented as mean ± SD. Statistical significance was determined using one-way ANOVA followed by Tukey’s multiple comparisons test. For (A), n = 7 representative images; for (B), n = 5 representative images per condition (*p < 0.05, **p < 0.01, ***p < 0.001,
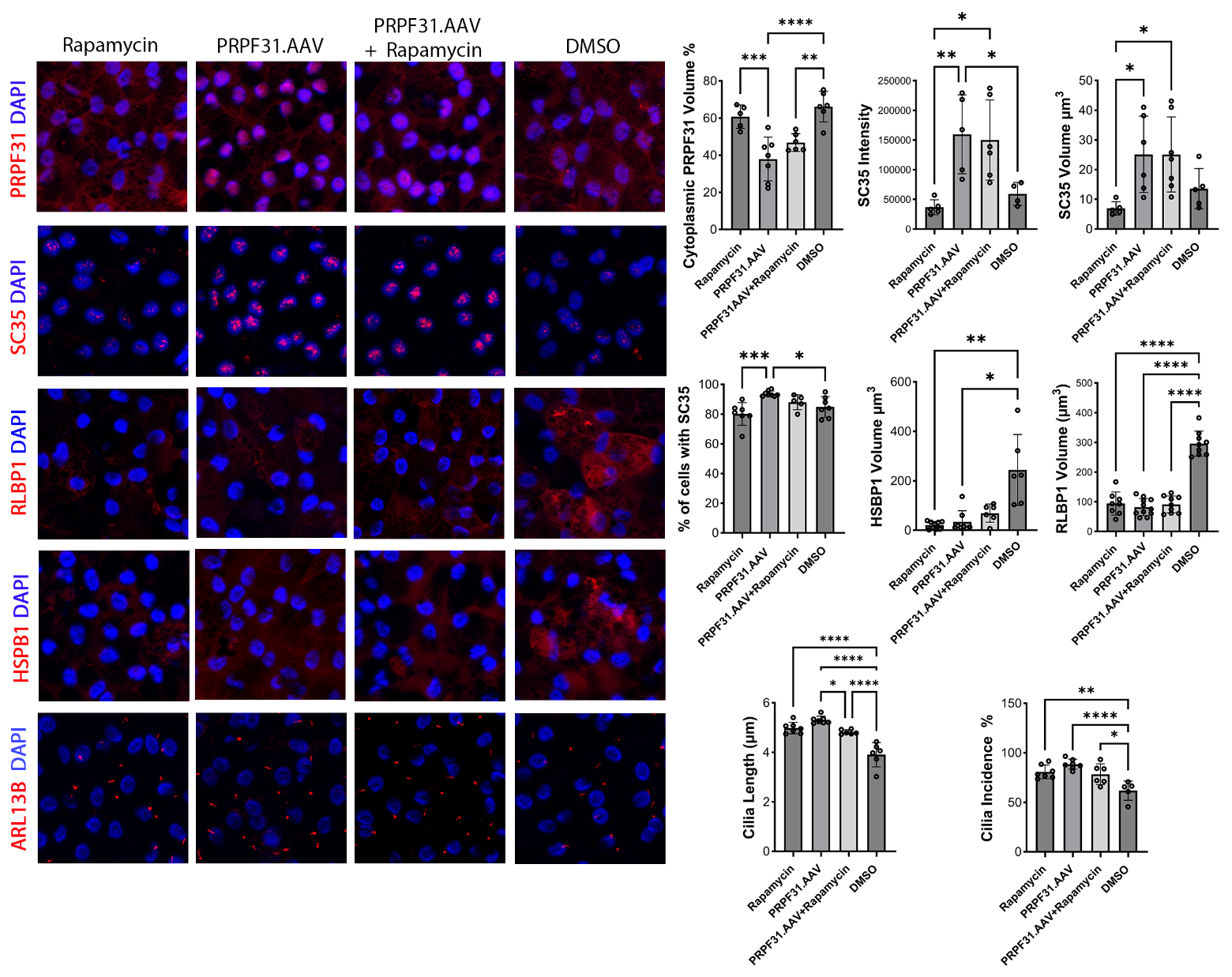
****p < 0.0001).

**Figure S4: Comparative Evaluation of PRPF31 Augmentation and Rapamycin-Induced Autophagy in Mature RP11-RPE Cells.**

Top Panel: Representative Immunofluorescence Imaging. Micrographs illustrate the therapeutic effects of *PRPF31*.AAV, Rapamycin, or combinatorial treatment on mature RP11-RPE. Markers include:

- Nuclear Homeostasis: PRPF31 (red) and SC35 (nuclear speckles; red).
- Proteotoxic Stress: RLBP1 and HSPB1 cytoplasmic aggregates (red).
- Ciliary Architecture: ARL13B (red).

Nuclei were counterstained with DAPI (blue). *PRPF31* augmentation successfully restored nuclear localisation, reorganised speckle morphology, cleared cytoplasmic aggregates, and lengthened primary cilia. In contrast, Rapamycin treatment alone only partially reduced the aggregate burden and provided no additive benefit when combined with *PRPF31*.AAV. Scale bars: 25 µm.

Right Panel: Quantitative Phenotypic Analysis. Comprehensive quantification of the rescue effects across all treatment groups. Metrics include PRPF31 volume (%), SC35 volume (µm3) and intensity, percentage of SC35+ cells (%), RLBP1 and HSPB1 aggregate volume (µm3), and cilia length (µm) and incidence (%).

Statistical Analysis: Data are presented as mean ± SD (n = 10 representative images per marker per condition). Statistical significance was determined by one-way ANOVA followed by Tukey’s multiple comparisons test.

**Supplementary table legends**

**Table S1:** Differential gene expression in RPE samples and retinal organoids treated with PRPF31.AVV versus GFP.AAV.

**Table S2:** Differential exon usage in retinal organoids and RPE cells treated with PRPF31.AAV versus GFP.AAV controls.

**Table S3:** Differential protein abundance between PRPF31.AAV and GFP.AAV RPE cells and associated GO analysis for biological processes and cellular components.

**Table S4:** Differential protein abundance between PRPF31.AAV and GFP.AAV retinal organoids and associated GO analysis for biological processes and cellular components.

**Table S5:** Summary of antibodies and primers used in this study.
